# The role of the epipeptide EpeX in defining competitive fitness in intra-species mixed isolate colony biofilms of *Bacillus subtilis*

**DOI:** 10.1101/2023.02.09.527868

**Authors:** Margarita Kalamara, James Abbott, Tetyana Sukhodub, Cait MacPhee, Nicola R. Stanley-Wall

**Affiliations:** Division of Molecular Microbiology, School of Life Sciences, University of Dundee, Dundee, DD5 4EH, UK; Data Analysis Group, Division of Computational Biology, School of Life Sciences, University of Dundee, Dundee, DD5 4EH, UK; National Biofilms Innovation Centre, School of Physics & Astronomy, University of Edinburgh, EH9 3FD Edinburgh, UK

**Keywords:** *Bacillus subtilis*, biofilm, kin discrimination, epipeptide, EpeX

## Abstract

Bacteria engage in competitive interactions with neighbours that can either be of the same or different species. Multiple mechanisms are deployed to ensure the desired outcome and one tactic commonly implemented is the production of specialised metabolites. The Gram-positive bacterium *Bacillus subtilis* uses specialised metabolites as part of its intraspecies competition determinants to differentiate between kin and non-kin isolates. It is, however, unknown if the collection of specialised metabolites defines competitive fitness when the two isolates start as a close, interwoven community that grows into a densely packed colony biofilm. Moreover, the identity of the most effective specialised metabolites has not been revealed. Here, we determine the competition outcomes that manifest when 21 environmental isolates of *B. subtilis* are individually co-incubated with the model isolate NCIB 3610 in a colony biofilm. We correlated these data with the suite of specialised metabolite biosynthesis clusters encoded by each isolate. We found that the *epeXEPAB* gene cluster correlated with a strong competitive phenotype. This cluster is responsible for producing the epipeptide EpeX. We demonstrated that EpeX is a competition determinant of *B. subtilis* in an otherwise isogenic context. When we competed the NCIB 3610 EpeX deficient strain against our suite of environmental isolates we found that the impact of EpeX in competition is isolate-specific, as only one of the 21 isolates showed increased survival when EpeX was lacking. Taken together, we have shown that EpeX is a competition determinant used by *B. subtilis* that impacts intra-species interactions in an isolate-specific manner.

## Introduction

Kin discrimination is the ability of individuals to discriminate against conspecific organisms based on phylogenetic relatedness, such that neighbouring cells with the closest phylogenetic relationship cooperate more than those that are more distantly related (1–3). It is believed that this behaviour has evolved to stabilise cooperation between isolates that share the same genes for cooperative traits (1–3) and exclude the more distantly related, thereby avoiding exploitation of communal secreted molecules by potential non-contributing neighbours (4, 5) and in doing so releasing nutrients and genetic material that can be scavenged (6).

The Gram-positive soil-dwelling bacterium *Bacillussubtilis* exhibits kin discrimination (7) and it is mediated by a combinatorial process where multiple genetic loci define the relationship between isolates. These loci primarily comprise genes encoding the production and response to antimicrobials and cell surface-modifying molecules (8). An experimental system that has been deployed to define the molecular basis for kin discrimination is the “swarm meeting assay”, where different isolates move (swarm, (9)) towards each other on a semi-solid surface from initially distinct inoculation positions. Kin strains are those that can intermingle, and non-kin strains are those that form a clearance zone between them (7).

Some of the molecules involved in kin discrimination are specialised metabolites, (also known as secondary metabolites) which are a diverse class of bioactive molecules (10). Specialised metabolites produced by *B. subtilis* that are involved in kin discrimination are sporulation-killing factor, subtilosin A, bacillaene, and sublancin 168 (8). The role of these specialised metabolites in intra-species interactions was strengthened by an examination of the growth inhibitory properties when a focal strain is grown in at a higher density on the surface of a lawn of the target strain. A correlation was drawn between isolates encoding different biosynthetic gene clusters and competition outcome (11). However, this correlation was not perfect, as in some cases isolates encoding the same suite of biosynthetic gene clusters could still inhibit the growth of each other (11). These analyses highlight the complexities in defining the outcome of intra-species interactions.

In this work, we were interested in understanding the molecules that govern the competitive dynamics of isolates growing within the same niche, a mixed isolate colony biofilm. Competitive fitness in a spatially constrained mixed community is known to be impacted by the spatial arrangement of the founding cells (12, 13)and by the presence of polymorphic toxins (14). However, knowledge surrounding the role that specialised metabolites play in shaping these interactions in mixed communities is lacking. Here, to address this knowledge gap, we set out to explore the relationship between the suite of specialised metabolite biosynthesis clusters (SMBC) encoded by 21 soil isolates of *B. subtilis* and the model isolate NCIB 3610 and their pairwise competitive fitness within colony biofilms. We obtained complete whole genome sequence data and detected the SMBC within all 22 genomes. We next correlated the presence of the accessory SMBCs with the competitive fitness of the isolates relative to the model isolate NCIB 3610. We identified that the SMBC whose presence most closely correlated with a strong competitive phenotype was the *epeXEPAB* cluster, which is responsible for the production of the epipeptide EpeX. We explored the role of EpeX in competitive fitness by constructing a deletion mutant of the biosynthetic cluster in the model isolate NCIB 3610. We found that, in an otherwise isogenic context, EpeX is an important determinant of competitive fitness, with the strain encoding the cluster occupying a higher proportion of the mixed community when compared with the NICB 3610 EpeX deficient mutant. When testing the generality of EpeX as an intra-species competition determinant, we identified one isolate within our suite of 21 isolates that exhibited increased survival when competed with the EpeX deficient strain of NCIB 3610 rather than the NCIB 3610 parental strain. Additionally, when exploring the role that EpeX has as a competition determinant in other isolates, we found that absence of the *epeXEPAB* cluster does not impact competitive fitness in two other soil isolates. In combination, our results reveal EpeX as a competition determinant of intra-species interactions but caution that the role it plays has an isolate-specific context.

## Methods

### Growth conditions and strains used

All strains used in this study are listed on Table 1. For routine growth of *Bacillus subtilis* and *Escherichia coli* strains, lysogeny broth (LB) liquid media was made using the following recipe: 1% (w/v) Bacto-peptone, 1% (w/v) NaCI, 0.5% (w/v) yeast extract. For solid plates, LB broth was supplemented with 1.5% (w/v) agar. The LB was sterilised by autoclaving. When necessary, LB media cultures and plates were supplemented with antibiotics which were used at the following concentrations for *B. subtilis*: 10 μg/ml kanamycin, 100 μg/ml spectinomycin and 5 μg/ml chloramphenicol. For growth of *E. coli* carrying plasmids of interest, the LB plates and liquid media were supplemented with 100 μg/ml of ampicillin, or 25 μg/ml chloramphenicol as required. Biofilm assays were conducted using MSgg (Minimal Salts glycerol glutamate) media. MSgg was made by first making a base medium, consisting of 5 mM potassium phosphate, 100 mM MOPS at pH 7.0, supplemented with 1.5% (w/v) agar. The media base was autoclaved and cooled to 55°C. The base medium was supplemented with 2 mM MgCl_2_, 700 μM CaCl_2_, 50 μM FeCl_3_, 50 μM MnCl_2_, 1 μM ZnCl_2_, 2 μM thiamine, 0.5% (v/v) glycerol and 0.5% (w/v) glutamic acid. A volume of 23 ml of MSgg melted media was added to each 9 cm diameter petri dish and the plates were solidified at room temperature. The surface of the solid plates was dried for 1 hour under a laminar flow cabinet prior to use in experiments.

**Table 1.** Strains used in this study

| Strain | Code Name <sup>a</sup> | Genotype <sup>b</sup> | Source <sup>c</sup> |
| --- | --- | --- | --- |
| NCIB 3610 |  | Wild type | B.G.S.C. |
| 168 |  | *trpC2* | B.G.S.C. |
| NRS6220 | NRS6103g | NRS6103 *amyE::Phy-spank-gfp mut2 (cml)* | pBL165 into NRS6103 |
| NRS6221 | NRS6105g | NRS6105 *amyE::Phy-spank-gfp mut2 (cml)* | pBL165 into NRS6105 |
| NRS6222 | NRS6153g | NRS6153 *amyE::Phy-spank-gfp mut2 (cml)* | pBL165 into NRS6153 |
| NRS6223 | NRS6096g | NRS6096 *amyE::Phy-spank-gfp mut2 (cml)* | pBL165 into NRS6096 |
| NRS6881 | NRS6085g | NRS6085 *amyE::Phy-spank-gfp mut2 (cml)* | pBL165 into NRS6085 |
| NRS6882 | NRS6099g | NRS6099 *amyE::Phy-spank-gfp mut2 (cml)* | pBL165 into NRS6099 |
| NRS6883 | NRS6107g | NRS6107 *amyE::Phy-spank-gfp mut2 (cml)* | pBL165 into NRS6107 |
| NRS6884 | NRS6116g | NRS6116 *amyE::Phy-spank-gfp mut2 (cml)* | pBL165 into NRS6116 |
| NRS6885 | NRS6118g | NRS6118 *amyE::Phy-spank-gfp mut2 (cml)* | pBL165 into NRS6118 |
| NRS6886 | NRS6121g | NRS6121 *amyE::Phy-spank-gfp mut2 (cml)* | pBL165 into NRS6121 |
| NRS6887 | NRS6127g | NRS6127 *amyE::Phy-spank-gfp mut2 (cml)* | pBL165 into NRS6127 |
| NRS6888 | NRS6128g | NRS6128 *amyE::Phy-spank-gfp mut2 (cml)* | pBL165 into NRS6128 |
| NRS6889 | NRS6132g | NRS6132 *amyE::Phy-spank-gfp mut2 (cml)* | pBL165 into NRS6132 |
| NRS6890 | NRS6145g | NRS6145 *amyE::Phy-spank-gfp mut2 (cml)* | pBL165 into NRS6145 |
| NRS6891 | NRS6160g | NRS6160 *amyE::Phy-spank-gfp mut2 (cml)* | pBL165 into NRS6160 |
| NRS6892 | NRS6181g | NRS6181 *amyE::Phy-spank-gfp mut2 (cml)* | pBL165 into NRS6181 |
| NRS6893 | NRS6183g | NRS6183 *amyE::Phy-spank-gfp mut2 (cml)* | pBL165 into NRS6183 |
| NRS6894 | NRS6186g | NRS6186 *amyE::Phy-spank-gfp mut2 (cml)* | pBL165 into NRS6186 |
| NRS6895 | NRS6187g | NRS6187 *amyE::Phy-spank-gfp mut2 (cml)* | pBL165 into NRS6187 |
| NRS6896 | NRS6190 | NRS6190 *amyE::Phy-spank-gfp mut2 (cml)* | pBL165 into NRS6190 |
| NRS6897 | NRS6202g | NRS6202 *amyE::Phy-spank-gfp mut2 (cml)* | pBL165 into NRS6202 |
| NRS6931 |  | 168 *amyE::Phy-spank-mTagBFP (spec)* | pNW2304 into 168 |
| NRS6932 | NCIB 3610b | NCIB 3610 *amyE::Phy-spank-mTagBFP (spec)* | NRS6931 SPP1 into NCIB 3610 |
| NRS6900 |  | 168 *amyE::Phy-spank-gfp mut2 (cml)* | pBL165 into 168 |
| NRS6942 | NCIB 3610g | NCIB 3610 *amyE::Phy-spank-gfp mut2 (cml)* | NRS6900 SPP1 into NCIB 3610 |
| NRS6934 | NRS6096b | NRS6096 *amyE::Phy-spank-mTagBFP (spec)* | pNW2304 into NRS6096 |
| NRS6935 | NRS6103b | NRS6103 *amyE::Phy-spank-mTagBFP (spec)* | pNW2304 into NRS6103 |
| NRS6936 | NRS6105b | NRS6105 *amyE::Phy-spank-mTagBFP (spec)* | pNW2304 into NRS6105 |
| NRS6937 | NRS6118b | NRS6118 *amyE::Phy-spank-mTagBFP (spec)* | pNW2304 into NRS6118 |
| NRS6938 | NRS6153b | NRS6153 *amyE::Phy-spank-mTagBFP (spec)* | pNW2304 into NRS6153 |
| NRS6943 | NRS6085b | NRS6085 *amyE::Phy-spank-mTagBFP (spec)* | pNW2304 into NRS6085 |
| NRS6944 | NRS6099b | NRS6099 *amyE::Phy-spank-mTagBFP (spec)* | pNW2304 into NRS6099 |
| NRS6945 | NRS6107b | NRS6107 *amyE::Phy-spank-mTagBFP (spec)* | pNW2304 into NRS6107 |
| NRS6946 | NRS6116b | NRS6116 *amyE::Phy-spank-mTagBFP (spec)* | pNW2304 into NRS6116 |
| NRS6947 | NRS6121b | NRS6121 *amyE::Phy-spank-mTagBFP (spec)* | pNW2304 into NRS6121 |
| NRS6948 | NRS6127b | NRS6127 *amyE::Phy-spank-mTagBFP (spec)* | pNW2304 into NRS6127 |
| NRS6949 | NRS6128b | NRS6128 *amyE::Phy-spank-mTagBFP (spec)* | pNW2304 into NRS6128 |
| NRS6950 | NRS6132b | NRS6132 *amyE::Phy-spank-mTagBFP (spec)* | pNW2304 into NRS6132 |
| NRS6951 | NRS6145b | NRS6145 *amyE::Phy-spank-mTagBFP (spec)* | pNW2304 into NRS6145 |
| NRS6952 | NRS6160b | NRS6160 *amyE::Phy-spank-mTagBFP (spec)* | pNW2304 into NRS6160 |
| NRS6953 | NRS6181b | NRS6181 *amyE::Phy-spank-mTagBFP (spec)* | pNW2304 into NRS6181 |
| NRS6954 | NRS6183b | NRS6183 *amyE::Phy-spank-mTagBFP (spec)* | pNW2304 into NRS6183 |
| NRS6955 | NRS6186b | NRS6186 *amyE::Phy-spank-mTagBFP (spec)* | pNW2304 into NRS6186 |
| NRS6956 | NRS6187b | NRS6187 *amyE::Phy-spank-mTagBFP (spec)* | pNW2304 into NRS687 |
| NRS6957 | NRS6190b | NRS6190 *amyE::Phy-spank-mTagBFP (spec)* | pNW2304 into NRS6190 |
| NRS6958 | NRS620b | NRS6202 *amyE::Phy-spank-mTagBFP (spec)* | pNW2304 into NRS6202 |
| NRS7253 |  | 168 *epeXEPAB::kan* | pNW2315 into 168 |
| NRS7259 | 3610g epe | NCIB 3610 *epeXEPAB::kan amyE::Phy-spank-gfp mut2 (cml)* | NRS7253 SPP1 into NRS6942 |
| NRS760 | 3610b epe | NCIB 3610 *epeXEPAB::kan amyE::Phy-spank-mTagBFP (spec)* | NRS7253 SPP1 into NRS6932 |
| NRS7390 | NRS6153g epe | NRS6153 *epeXEPAB::kan amyE::Phy-spank-gfp mut2 (cml)* | pNW2315 into NRS6222 |
| NRS7391 | NRS6202g epe | NRS6202 *epeXEPAB::kan amyE::Phy-spank-gfp mut2 (cml)* | pNW2315 into NRS6897 |
| NRS7392 | NRS6153b epe | NRS6153 *epeXEPAB::kan amyE::Phy-spank-mTagBFP (spec)* | pNW2319 into NRS6938 |
| NRS7393 | NRS6202b epe | NRS6202 *epeXEPAB::kan amyE::Phy-spank-mTagBFP (spec)* | pNW2319 into NRS7201 |

**a** The naming given to strains in figures and figure legends is indicated
**b** The abbreviation “spec” indicates spectinomycin resistance; “cml” indicates chloramphenicol resistance and “kan” kanamycin resistance.
**c** The method of strain construction is indicated with either the plasmid (pNW) or donor strain phage (SPP1) inserted into the parental strain.

### Strain construction

The strain used for storing of plasmids for cloning was *Escherichia coli* strain MC1061 [F’ *laclQ lacZM15 Tn*10 (*tet*)]. For making mutations in the NCIB 3610 background, as this strain is not genetically competent, plasmids were first transformed into the laboratory strain 168 through using standard protocol (15). The modified region was subsequently inserted and integrated into the NCIB 3610 genome via SPP1 phage transduction (16). For genetically competent soil isolates of *B. subtilis*, the plasmids were transformed directly into the isolate of interest as previously described (17) with the adaptations described in (18).

The *epeXEPAB* deletions in the *B. subtilis* isolates were constructed by homologous recombination and insertion of a kanamycin resistance cassette in the native locus, using plasmid pNW2315. For construction of pNW2315 the required fragment was synthesised by GenScript and inserted into the pCC1 vector. The construct sequence can be found in Table S1. Strains with the *epeXEPAB* deletion were verified by using the primers NRS2812 (5’ GTCTCGTATAATCTCTCACTTTCCC 3’) and NRS3311 (5’ AGTAAGTGGCTTTATTGATCTTGGG 3’).

For construction of the mTagBFP and GFP-expressing isolates, plasmids pNW2304 (12) and pBL165 (19) were used respectively. Both plasmids are designed to facilitate the integration of the genes encoding the fluorescent proteins and antibiotic resistance cassettes into the *amyE* locus. Resulting colonies were therefore screened using a potato starch assay to assess loss of amylase activity (20) and expression of the appropriate fluorescent protein.

### Biofilm co-culture assays

The mixed biofilm assays were set up as previously described (12). Cultures of the individual strains to be used were set up in 5 ml of LB and incubated at 37°C with agitation overnight. The following morning, day cultures were set up by inoculating 3 ml of LB with 200 μl of the overnight cultures. The day cultures were incubated at 37°C with agitation. The growth of the cultures was monitored, and all cultures were normalised to an OD_600_ of 1. After normalisation, cultures were mixed at a 1:1 ratio as required. 5 μl drops of the culture mixtures were spotted onto MSgg agar plates and 5 μl drops of the individual normalised cultures were included in the assays as controls. The plates were incubated at 30°C and images were taken after 24,48 and 72 hours as required. Fluorescence imaging was performed using a Leica fluorescence stereoscope (M205FCA) with a 0.5 × 0.2 NA objective. Imaging files were imported to OMERO (21).

### Image analysis

Relative strain densities of GFP and mTagBFP-expressing cells in mixed biofilm assays were determined by analysing fluorescent imaging data. This was done using a macro which was kindly produced by Dr. Graeme Ball at the Dundee Imaging Facility. Fiji/lmageJ (11, 12) was used to run the macro as previously described in our publication (12). Figures were constructed using GraphPad prism 7.

### Enhanced whole genome sequencing

Enhanced whole genome sequencing was performed by MicrobesNG. This required a combination of Illumina short-read data acquisition and nanopore sequencing for long-read data. For the preparation of samples in the lab, a single colony of each strain to be sequenced was resuspended in 200 μl of sterile PBS buffer and 100 μl of this was used to inoculate 300 ml of LB broth. The remaining 100 μl was streaked on an LB agar plate, which was incubated at 37°C overnight. The 300 ml culture was incubated at 16°C with shaking overnight. The following morning, the culture was incubated at 37°C with shaking and the OD_600_ was monitored. When cultures had reached an OD_600_ value of between 0.5 and 0.8, they were centrifuged at 3,750 rpm for 10 minutes. The supernatant was removed, and the pellets were resuspended in a tube with a cryopreservative (Microbank™, Pro-Lab Diagnostics UK, United Kingdom) or with DNA/RNA Shield (Zymo Research, USA) following MicrobesNG strain submission procedures. The weight of the pellet required for *B. subtilis* submission was at least 1 gram, so all samples were grown in large enough volumes to exceed 1 gram of pelleted cells. The spread plate set up at the same time as the culture was used for quality assessment, to ensure no contamination had occurred. The samples were sent to the MicrobesNG facilities. There, for DNA extraction, 5 to 45 μl of the suspension was lysed with 120 μl of TE buffer containing lysozyme (final concentration 0.1 mg/mL)and RNase A (ITW Reagents, Barcelona, Spain) (final concentration 0.1 mg/mL), incubated for 25 min at 37°C. Proteinase K (VWR Chemicals, Ohio, USA) (final concentration 0.1 mg/mL) and SDS (Sigma-Aldrich, Missouri, USA) (final concentration 0.5% v/v) were added and incubated for 5 min at 65°C. Genomic DNA was purified using an equal volume of SPRI beads and resuspended in EB buffer (Qiagen, Germany). DNA was quantified with the Quant-iT dsDNA HS kit (ThermoFisher Scientific) assay in an Eppendorf AF2200 plate reader (Eppendorf UK Ltd, United Kingdom). For Illumina sequencing, genomic DNA libraries were prepared using the Nextera XT Library Prep Kit (Illumina, San Diego, USA) following the manufacturer’s protocol with the following modifications: input DNA was increased 2-fold, and PCR elongation time was increased to 45 s. DNA quantification and library preparation were carried out on a Hamilton Microlab STAR automated liquid handling system (Hamilton Bonaduz AG, Switzerland). Pooled libraries were quantified using the Kapa Biosystems Library Quantification Kit for Illumina. Libraries were sequenced using Illumina sequencers (HiSeq/NovaSeq) using a 250bp paired end protocol. Long read genomic DNA libraries were prepared with Oxford Nanopore SQK-RBK004 kit and/or SQK-LSK109 kit with Native Barcoding EXP-NBD104/114 (ONT, United Kingdom) using 400-500ng of HMW DNA. Barcoded samples were pooled together into a single sequencing library and loaded in a FLO-MIN 106 (R.9.4.1) flow cell in a GridlON (ONT, United Kingdom).

### Genome Assembly

Illumina reads were adapter trimmed using Trimmomatic 0.30 with a sliding window quality cutoff of Q15 (22). An initial nanopore-only genome assembly was carried out using Flye 2.9.1 (23) with the ‘nano-raw’ model, and the resulting contigs used in conjunction with the Illumina reads with Unicycler v0.5.0 (24) using ‘bold’ mode to produce a final assembly. The resulting contigs were annotated using bakta 1.40 (database version 3.1) (25). Examination of the assembly graphs allowed putative plasmid sequences to be identified in cases where short, circular molecules were evident which were not integrated into the chromosomal sequence. Raw sequence reads and annotated assemblies can be found under European Nucleotide Archive Project PRJEB43128.

### Phylogenetic tree construction

The nucleotide sequences of *gyrA, rpoB, dnaJ* and *recA* were extracted from the short read data (which can be found in our previous publication (18)) using Artemis (26) and concatenated. The same sequences for the reference strain *B. subtilis* NCIB 3610 (Genbank accession number GCA_002055965.1) were retrieved from NCBI, concatenated, and included in the analysis. The sequences were aligned in Jalview (27) by MAFFT using the G-INS-I algorithm and MEGA7 software (28) was used to constructa maximum likelihood phylogenetic tree with 100 bootstrap repeats as previously described (18).

### Pangenome analysis

A pangenome analysis of all environmental isolates included in this work, the model isolate NCIB 3610 and other publicly available genome sequences of *B. subtilis* isolates was constructed using Roary version 3.13.0 with default parameters. The draft genome assemblies were used as the input. The pangenome figure was produced using the roary_plots.py macro and further annotated in Adobe Illustrator (https://adobe.com/products/illustrator)

### Command line blast

To explore the presence and distribution of the genes within the *epeXEPAB* cluster, command line blast was used to create a nucleotide database using the whole genomes of NCIB 3610 and the 21 genetically competent isolates in our collection. The database was then used to perform nucleotide blast searches of the *epe* genes. The outcome of the analysis and locations of genes of interest were used to manually extract the sequences of interest. The sequences were aligned and exported as image files in Jalview (27) to explore the diversity in the coding sequences where required.

### antiSMASH

To determine the secondary metabolite biosynthesis clusters encoded by each isolate, antiSMASH version 6.0 was used (29). Enhanced whole genome sequence assemblies were submitted to the server and run with default settings. Genbank files of all secondary metabolite biosynthesis clusters encoded by all isolates were retained.

### Clinker

Clinker version 0.0.20 was used with default settings to visualise the secondary metabolite biosynthesis clusters identified by antiSMASH. The GenBank files of the clusters downloaded from antiSMASH were used as an input for clinker to produce figures. The figures were modified using Adobe Illustrator (https://adobe.com/products/illustrator).

## Results

### Mixed biofilm intra-species competition and phylogenetic relatedness

We examined the competitive outcome of the interaction between 21 genetically competent *B. subtilis* soil isolates and the model isolate NCIB 3610 in the context of a mixed isolate colony biofilm. In each case, we competed a variant of NCIB 3610 that constitutively expresses mTagBFP against the GFP expressing variants of the soil isolates. We also included an NCIB 3610 isogenic mix as a control. We imaged the colony biofilms after 24,48 and 72 hours of incubation at 30°C (Figure S1). We quantified the proportion of GFP-expressing cells in the mixed biofilm using image analysis methods (12) (Figure 1A). Analysis of the NCIB 3610 isogenic control revealed that the GFP variant typically comprises approximately 60% of the community. As a 1:1 ratio between GFP and mTagBFP variants of NCIB 3610 is expected, the slight under representation of the mTagBFP variant is perhaps due to differences in fitness associated with the different fluorescent proteins (Figure 1B). The underrepresentation of the strain carrying mTagBFP is consistent with our previous observations and did not preclude us from defining the relationships between the isolates (12).

**Figure 1:**
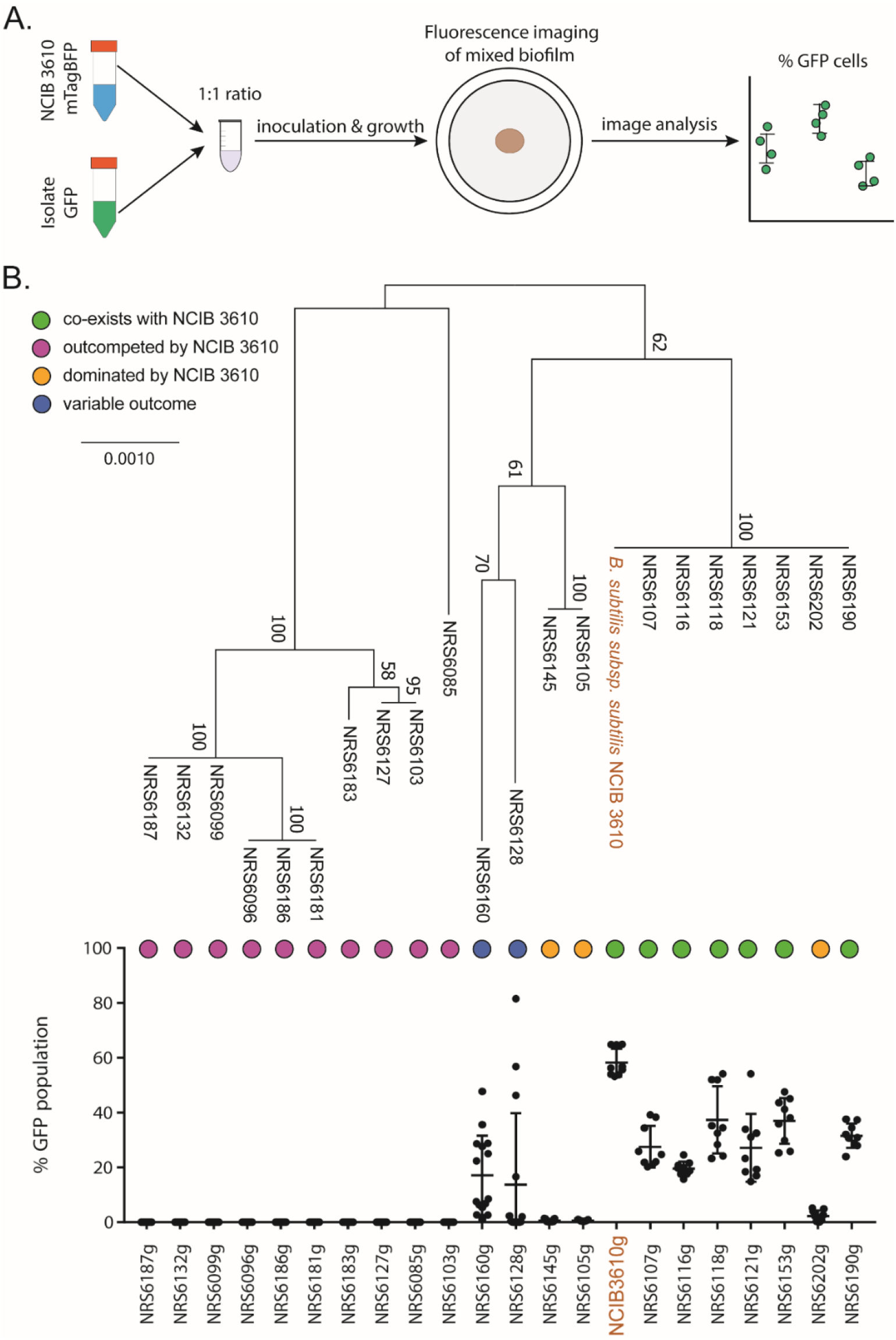
Mixed biofilm intra-species competition outcomes of *B. subtilis* isolates against the model NCIB 3610. (A) Schematic representation of mixed biofilm setup. (B) Maximum likleihood phylogenetic tree based on the concatenated sequences of housekeeping genes *gyrA, rpoB, dnaJ, recA* shown alongside the competition outcomes of mixed biofilms of NCIB 3610 co-incubated with each of the 21 environmental isolates of *B. subtilis* used in this work. The presented values are the % of the community of GFP expressing soil isolates, quantified using image analysis. The nine data points presented for each isolate represent three biological repeats and three technical repeats. The error bars represent the standard deviation of the mean.

The outcome of competition between the pairs of isolates shows that NCIB 3610 is a strong competitor that outcompetes most soil isolates from the 24h time point (Figure 1B, Figure S1). It is also evident that, for isolate pairs where co-existence is observed at the 24h timepoint, the proportion of soil isolate in the community decreases overtime (Figure S1, Figure S2). Based on the outcome of their interaction with NCIB 3610 after 24 hours of co-incubation, we defined the isolates in our collection as “outcompeted” (those that took up 0% of the community), “dominated” (those that took up 0-5% of the community), “co-existing” (those that took up more than 5%) and “variable” (those that in some rounds were dominated and in others co-existed) (Figure 1B, Figure S2A) using custom thresholds.

*B. subtilis* intra-species interactions have primarily been studied in the context of kin discrimination, which is defined as the differential treatment of conspecific isolates based on phylogenetic relationship (6–8, 30–32). Therefore, we correlated the outcome of the mixed biofilm screens with a maximum likelihood tree based on the concatenated nucleotide sequences of four housekeeping genes (*gyrA, rpoB, recA, dnaJ*). Our results show there is a correlation between the ability of isolates to co-exist with NCIB 3610 and how related the isolates are. All isolates that co-exist with NCIB 3610 are in the same phylogenetic group. Only one isolate of this group (namely NRS6202) fell within the class of isolates that were dominated by NCIB3610. The remaining two isolates that were in the “dominated” group, along with the two isolates that show “variable” results, are more distantly related to NCIB 3610. All isolates that are “outcompeted” by NCIB 3610 form the most distantly related phylogenetic groups (Figure 1B). This analysis indicates that the outcome of the interactions between our isolates are broadly consistent with the concept of kin discrimination.

### Pangenome analysis of soil isolates of *B. subtilis*

The 21 isolates of *B. subtilis* used in this work have been isolated from soil samples in Scotland (18). To explore the genomic diversity of these isolates, we used short read sequence data (18) and performed a pangenome analysis using Roary (33). We included all the isolates in our collection (18) alongside other selected publicly accessible genomes to provide coverages of other geographic locations and isolation sources. The analysis shows that there is a large diversity in the accessory genes found within the isolates examined. Additionally, the phylogenetic distribution of the isolates in our collection is varied with isolates positioned within different clades (Figure 2). Importantly, the analysis shows that the isolates in our collection, while sampled locally, provide a good representation of the diversity found among more widely sampled *B. subtilis* isolates. To facilitate further bioinformatic analysis we acquired the enhanced whole genome sequences for the isolates (MicrobesNG, Birmingham, United Kingdom). After receiving the illumina reads and long read data, the genomes were quality assessed and re-assembled to incorporate our initial Illumina data (18) and consequently increase coverage (Table S2) (ENA Project PRJEB43128).

**Figure 2:**
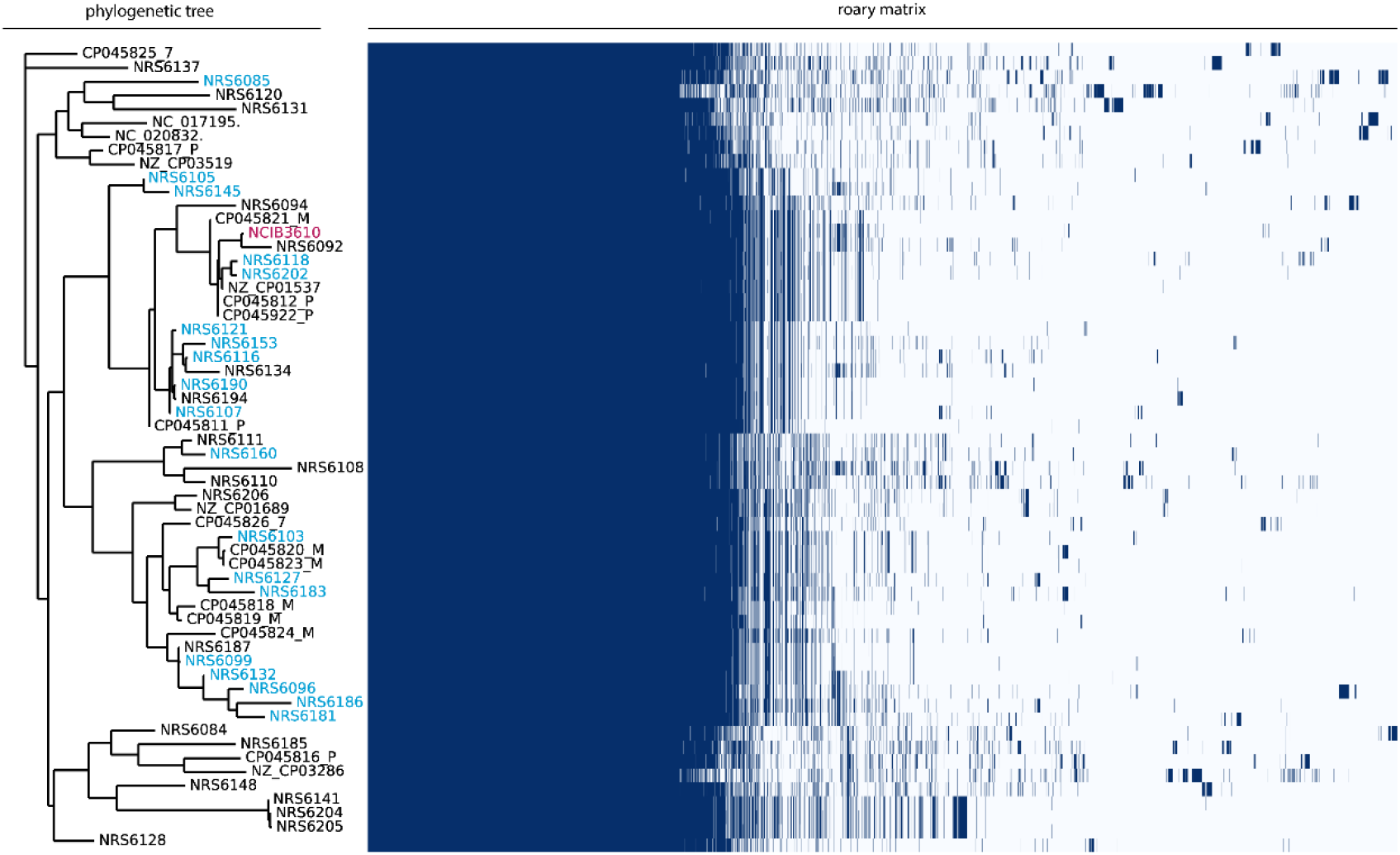
Pangenome analysis and phylogeny of *B. subtilis* isolates. The names of genetically competent soil isolates from the NSW laboratory are coloured in blue on the phylogenetic tree shown on the left. Non-competent isolates in the NSW lab collection and publicly accessible genomes from diverse sources are coloured in black. The model isolate NCIB 3610 is shown in pink. The roary matrix shows the presence (blue) and absence (white) of genes in each isolate.

### Exploring the specialised metabolite biosynthesis clusters encoded by the isolates in our collection

To uncover the specialised metabolite biosynthesis clusters (SMBC) encoded by each of the isolates in our collection we used antiSMASH version 6.0 (29), a tool designed for mining bacterial genomes and detecting such clusters. We correlated the presence of SMBCs that have a known antimicrobial function with the competitive phenotype of our isolates (Figure 3). In some cases, sequence variations and truncations were found in SMBCs for a small subset of isolates (Figure 3, Figure S3). The core clusters, a version of which was present in all isolates in our collection, are those required for the biosynthesis of bacillaene (34), plipastatin (35), bacillibactin (36), surfactin (37), subtilosin A (38) and bacilysin (39). One hypothesis is that the differential regulation of the core clusters could explain the competition outcome. However, here we focused on clusters that were not contained in all the genomes which produce metabolites with known antimicrobial properties, as we considered these likely to be involved in intra-species competition. The variable clusters encoded in our collection of *B. subtilis* isolates were those responsible for producing subtilomycin (40), sporulation killing factor (41–43), epipeptide (44, 45) and sublancin 168 (46, 47) (Figure 3, Table S3). Of these clusters we chose to further investigate the operon encoding for the epipeptide, as the presence of this cluster most closely correlated with a strong competitive phenotype (Figure 3). Only NCIB 3610 and isolates that could survive in the presence of NCIB 3610 encoded the entire cluster.

**Figure 3:**
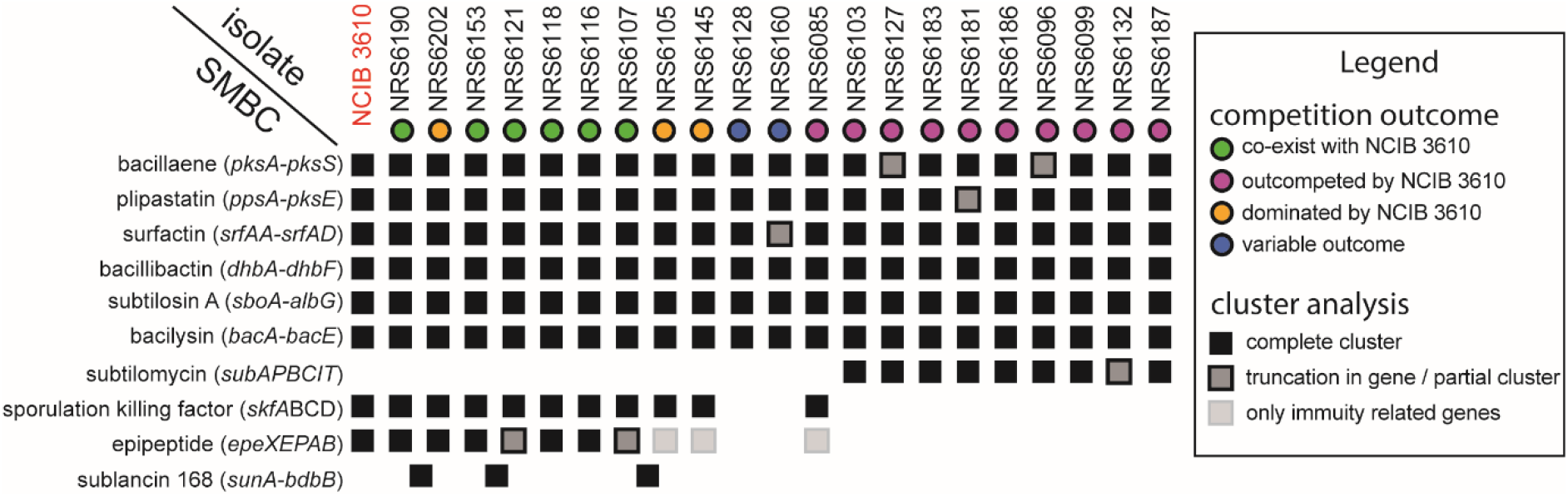
Secondary metabolite biosynthesis clusters and competitive fitness of soil isolates of *B. subtilis*. The specialised metabolites on the left-hand side represent the molecules encoded by each cluster identified by antiSMASH (29). The NCIB 3610 and numbers followed by “NRS” at the top represent different isolates used in this study. The outcomes of competitions in biofilms are indicated by coloured circles. This data is presented in Figure 1B and are as shown in the legend. The coloured squares show the presence and any variations in the encoded clusters and what they represent is shown in the legend.

### EpeX is a potential competition determinant

The *epe* cluster of *B. subtilis* NCIB 3610 consists of *epeX, epeE, epeP, epeA* and *epeB* (Figure 4A). The variants of the cluster found within our isolate collection are presented (Figure 4A) and full details are provided (Table S4, Figure S4, Figure S5, Figure S6). EpeX has a toxic effect on the cell envelope of *B. subtilis* (45, 48). It is made as pre-pro-peptide in the cytoplasm that is processed by the radical-S-adenosyl-L-methionine (SAM) epimerase EpeE, which converts the L-valine and L-isoleucine of EpeX into their D-configured counterparts generating pre-EpeX (49). Pre-EpeX is further exported and cleaved, and based on the genomic arrangement, it is predicted that this is mediated by EpeP, a membrane anchored signal peptidase (44). Finally, EpeAB form an ABC transporter that confers partial resistance to the intrinsically produced EpeX and is involved in autoimmunity (48) (Figure 4B). The EpeX peptide triggers the activation of the LiaRS-dependent cell envelope stress response, and LiaH (phage heat shock protein) and Lial (membrane anchor) are additional major resistance determinants against the antimicrobial peptide. Consistent with the cell envelope stress reponse being involved in immunity against the epipeptide, the mode of action of EpeX is membrane depolarization which causes permeabilization of the mebrane (45). This makes EpeX a likely candidate for a role in intra-species interactions and kin discrimination.

**Figure 4:**
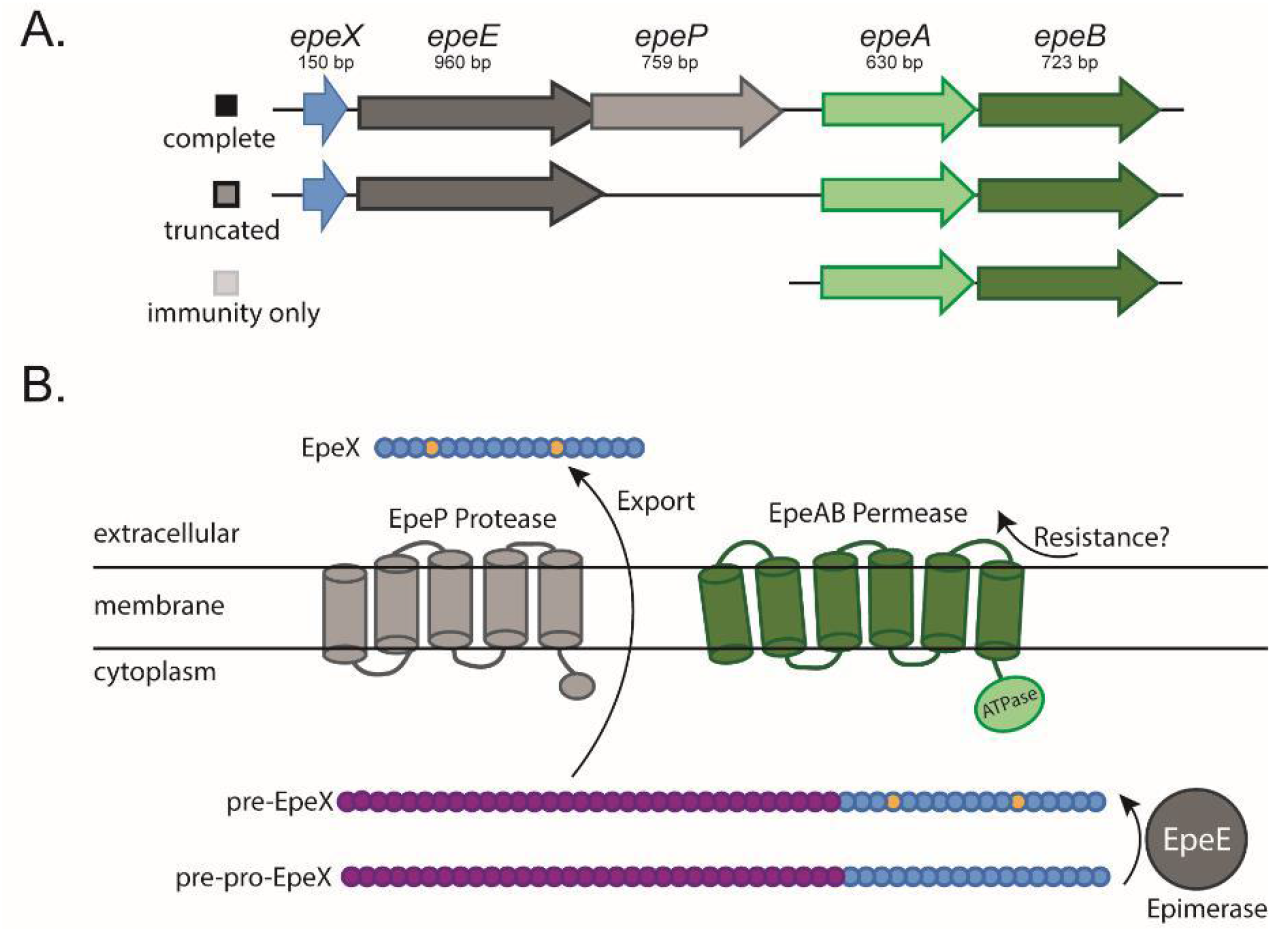
EpeX as a potential competition determinant of intra-species interactions. **(A)** Schematic representation of the variants of the *epeXEPAB* found in the genomes of the isolates used in this work. The coloured boxes next to each cluster schematic identify the cluster variant; (B) schematic representation of the components and function of EpeXEPAB. Amino acids coloured in yellow for the pre-EpeX indicate amino acids epimerised by EpeE prior to being cleaved and is presumably further processed and exported from the cell. The processing and export are thought to be mediated by the EpeP protease to generate the final form, EpeX. The EpeAB permease is believed to be involved in immunity against EpeX. The schematic has been adapted from (44).

### Absence of the *epeXEPAB* cluster impacts competition against an otherwise isogenic strain

To investigate if the *epeXEPAB* cluster has a role in shaping intra-species interactions in the context of mixed isolate colony biofilm, we constructed a variant of NCIB 3610 that lacks the entire *epeXEPAB* cluster. We tested the competitiveness of this mutant against NCIB 3610 in mixed isolate colony biofilms. From the single isolate controls, it is apparent that, at least on a macroscopic level, colony morphology is not impacted by an absence of the *epeXEPAB* cluster (Figure 5A). To determine the outcome of the competition between the strains in the mixed colony biofilms, we again used image analysis to quantify the proportion of GFP and mTagBFP expressing cells in the community that developed. Our results shown that the *epeXEPAB* mutant of NCIB 3610 is less successful than the wild type, as the proportion of the community it occupied is significantly lower than that taken up by the wild type in the isogenic control sample (Figure 5B and C). These data show that the *epeXEPAB* cluster is a determinant of the competition outcome in an otherwise isogenic biofilm co-culture. Lack of this cluster decreases the competitive strength of *B. subtills* NCIB 3610.

**Figure 5:**
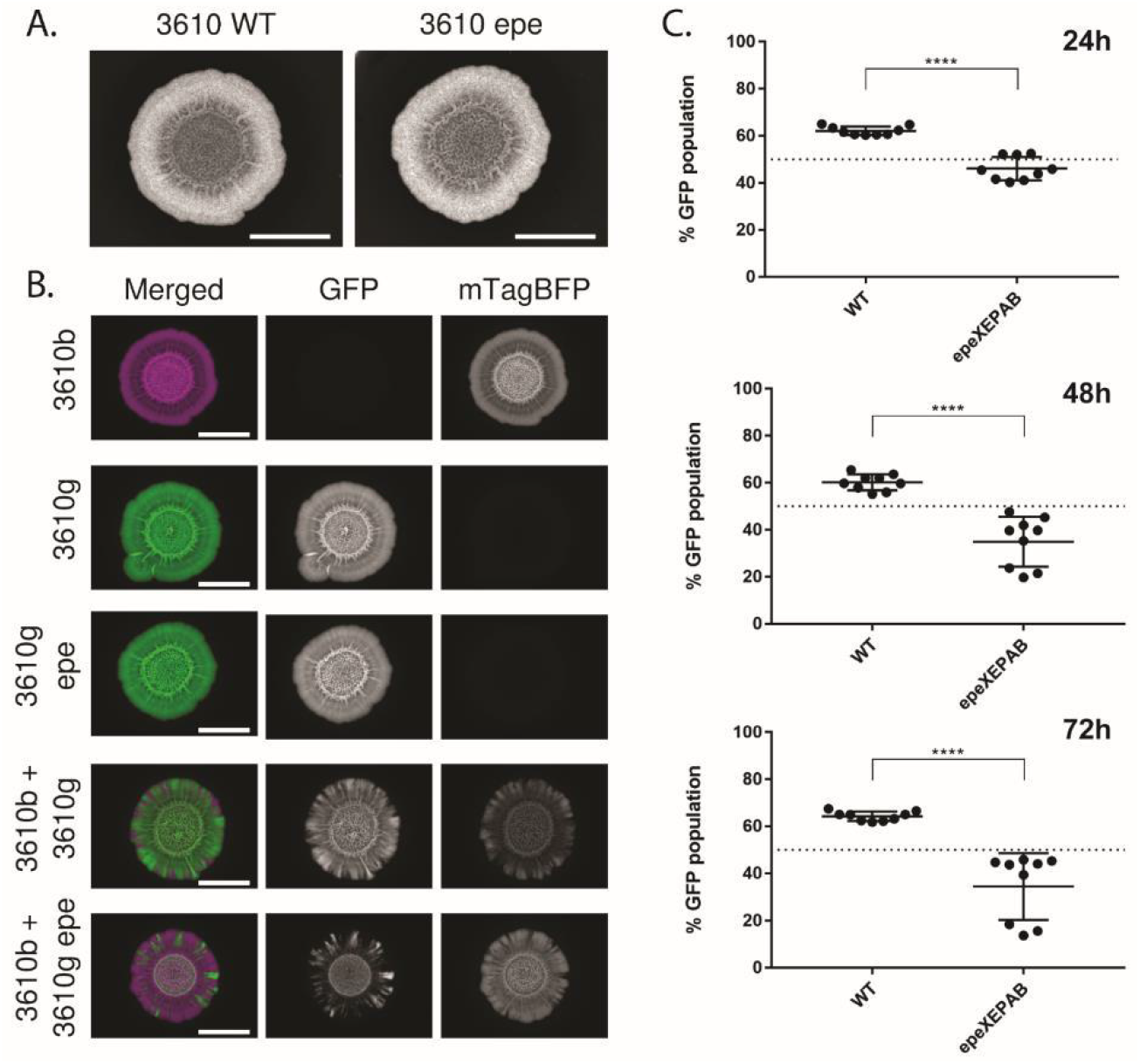
Competition assay outcome between NCIB 3610 wild type and *epeXEPAB* mutants. (A) Representative images of single strain biofilms of the wild type (“WT”) and *epeXEPAB* (“epe”) mutant of NCIB 3610 (“3610”) grown on MSgg media for 48 hours at 30°C. The scale bars represent 0.5 cm. (B) representative images of biofilms growth for 48 hours at 30°C on MSgg agar. “3610” is the model isolate NCIB 3610. Strain names followed by “b” represent strains constitutively expressing mTagBFP, false coloured in magenta and names followed by “g” represent strains constitutively expressing GFP and are false coloured in green, “epe” represents deletion of the *epeXEPAB* operon. “3610b” and “3610g epe” are images of the same biofilms as those shown in (A). The scale bars represent 0.5 cm. (C) Competition results of NCIB 3610 wild type expressing mTagBFP (NRS6932) against GFP-expressing wild type (NRS6942) or *epeXEPAB* mutant (NRS7259) of NCIB 3610 as indicated after 24, 48 and 72 hours of co-incubation on biofilm inducing media plates as indicated. The presented values are the % of the community of GFP expressing strain, quantified using image analysis. Each individual data point presented for each isolate represent one of two or three technical replicates for the three biological repeats performed. The error bars represent the standard deviation of the mean. The asterisks represent statistical significance with a p value of ≤ 0.0001 between the two populations as calculated using an unpaired t test.

### A limited role for EpeX as an intraspecies competition determinant

Next, we explored how NCIB 3610 lacking the *epeXEPAB* cluster competed when mixed with the 21 soil isolates in our collection. We hypothesised that if EpeX is a competition determinant of intra-species interactions, then the lack of *epeXEPAB* would reduce the competitive fitness of NCIB 3610. This would allow for a) under representation of the NCIB 3610 *epeXEPAB* strain in cases where co-existence was achieved with the wild type, and/or *b*) isolates that are outcompeted or dominated by the wild NCIB 3610 managing to achieve some level of co-existence with the *epeXEPAB* mutant. We used an mTagBFP-expressing variant of NCIB 3610 *epeXEPAB* as a reference strain, competing it against our suite of GFP-expressing isolates, and overlayed the data from this screen with the data obtained from the screen of all isolates against the wild type NCIB 3610 (recall Figure 1B). Our results show that for most of the isolates, the loss of the *epeXEPAB* cluster in NCIB 3610 has no impact on the outcome of the pairwise competition (Figure 6A). The only isolate that takes up a larger portion of the community when mixed with the *epeXEPAB* mutant versus the wild type of NCIB 3610 is isolate NRS6153. To explore this relationship more closely, we further analysed the data and found that there is a statistically significant difference between the portion of the community taken up by NRS6153 when mixed with the two variants of NCIB 3610 (Figure 6B). However, deletion of the *epeXEPAB* cluster in NRS6153 did not impact competition with its otherwise isogenic parental strain (Figure S7A). This was also the case for NRS6202 (Figure S7B). Collectively, our data uncover a role for EpeX as a competition determinant of *B. subtilis* intra-species interactions but reveal that the impact that EpeX has varies greatly depending on the competing isolate.

**Figure 6:**
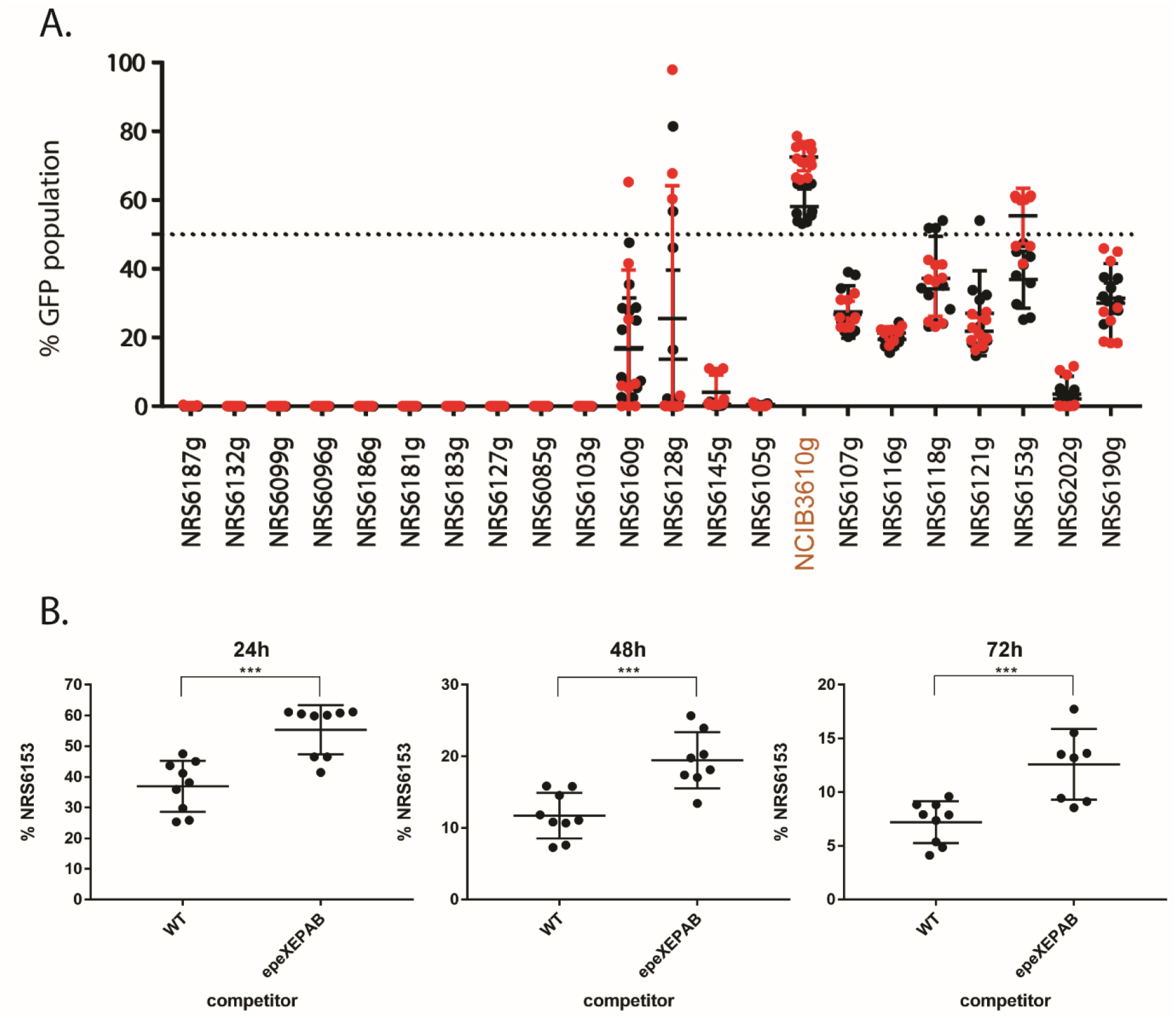
comparison of mixed biofilm outcomes using NCIB 3610 wild type and *epeXEPAB* as references. (A) Competition results of NCIB 3610 wild type (WT) expressing mTagBFP (NRS6932, black data points) or NCIB 3610 *epeXEPAB* expressing mTagBFP (NRS7260, red data points) against GFP-expressing soil isolates at 24 hours of co-incubation on biofilm inducing media plates as indicated. The presented values are the % of the community of GFP expressing soil isolates, quantified using image analysis. Each individual data point presented for each isolate represent one of two or three technical replicates for three biological repeats with each reference strain as indicated. The error bars represent the standard deviation of the mean (B) Competition results of GFP-expressing NRS6153 (NRS622) against *mTagBFP*-expressing wild type (NRS6932) or *epeXEPAB* mutants (NRS7260) of NCIB 3610 after 24, 48 and 72 hours of co-incubation on biofilm inducing media plates as indicated. The presented values are the % of the community of GFP expressing strain (NRS6153), quantified using image analysis. Each individual data point presented for each isolate represent one of two or three technical replicates for the three biological repeats performed. The error bars represent the standard deviation of the mean. The asterisks represent statistical significance with a p value of ≤ 0.001 between the two populations as calculated using an unpaired t test.

## Discussion

In this study, we combined bioinformatic analysis and physiological experiments to identify a new competition determinant of *B. subtilis* intra-species interactions that is active within a spatially confined colony biofilm. By assessing genome data alongside the outcomes of pair-wise competitions of 21 soil isolates challenged against NCIB 3610, we found a correlation between isolates encoding the cluster responsible for producing the epipeptide EpeX and competitive fitness. We hypothesised thatthis cluster was (in part) responsible for increasing the competitive fitness of isolates in a conspecific competition setting. To test our hypothesis, we deleted this cluster in NCIB 3610 and performed competitions of the mutant against both the wild type NCIB 3610 and our collection of soil isolates. We found that lack of the cluster responsible for EpeX production led to a decrease in competitive fitness in an otherwise isogenic context for NCIB 3610. When the variant of NCIB 3610 lacking the *epe* cluster was competed against the rest of the isolates in the strain collection it displayed the same competitive strength as the parental isolate for 20 of the 21 isolates. The exception was isolate NRS6153 where it occupied a significantly larger portion of the mature colony biofilm community when mixed against the *epeXEPAB* mutant compared with its pairing with the wild type NCIB 3610. Additionally, looking beyond the model isolate NCIB 3610, when we deleted the *epeXEPAB* cluster in isolates NRS6153 and NRS6202, no impact on competitive fitness was observed.

The identification of EpeX as a competition determinant within the spatially confined colony biofilm is consistent with the cluster being expressed during biofilm formation. If the production of EpeX did not coincide with the conditions used, no impact of removing the molecule would be observed. Activity of the epipeptide within a colony biofilm is also consistent with what is known about the expression profile of the *epe* operon. A critical regulator of biofilm matrix production and sporulation, Spo0A (50) relieves the repression of *epe* transcription via AbrB to allow EpeX to be produced (44). The reason why there is an isolate specific response to the presence of the epipeptide between different isolates remains to be explored. One possible explanation is the fact that immunity against EpeX is not straight-forward and is largely achieved through activation of the broad cell envelope stress response, orchestrated by the LiaRS two components system (45,48). Therefore, potential differences in the timing and combination of cell wall targeting competition determinants under the conditions tested could result in various levels of susceptibility of target cells to EpeX and the observed differences in the impact that this molecule has on competition. One way to explore how NCIB 3610 induces LiaRS response in different isolates could be using transcriptional reporter fusions with the promoter of the LiaRS system in both isolates that are impacted by EpeX and those that are not.

### Overarching Conclusion

Specialised metabolites are important determinants of social interactions among bacteria. While it is known that some specialised metabolites impact kin discrimination in the context of swarm meeting assays (8), it was unknown if and how different specialised metabolites affect the competitive strength of an isolate against conspecific isolates in a mixed biofilm. As biofilm formation is a very different physiological state to swarming (51) it is unknown if the molecules that affect mixing of swarms will be the same as those impacting competition in a biofilm setting. Additionally, the swarm meeting assays used previously to define the molecular determinants of kin discrimination (8) do not give any information about the competitive fitness of individual isolates, but rather just determine whether two strains can share a niche or not. In this work we addressed some of these knowledge gaps and revealed EpeX as a novel competition determinant, albeit with limited influence among other isolates.

## Acknowledgements

Work in the NSW and OEM laboratories was funded by the Biotechnology and Biological Science Research Council (BBSRC) [BB/P001335/1, BB/R012415/1]. M.K. was supported by a Biotechnology and Biological Sciences Research Council studentship [BB/M010996/1]. We are grateful to Joana Moreira Carneiro for her help with experimental work.

## Conflicts of Interests

There are no conflicts of interest to report.

## CRediT

| Term | Definition | Initials |
| --- | --- | --- |
| Conceptualization | Ideas; formulation or evolution of overarching research goals and aims | MK, NSW, CEM |
| Methodology | Development or design of methodology; creation of models | MK, NSW, CEM, JA |
| Software | Programming, software development; designing computer programs; implementation of the computer code and supporting algorithms; testing of existing code components | JA |
| Validation | Verification, whether as a part of the activity or separate, of the overall replication/ reproducibility of results/experiments and other research outputs | N/A |
| Formal analysis | Application of statistical, mathematical, computational, or other formal techniques to analyze or synthesize study data | MK, NSW |
| Investigation | Conducting a research and investigation process, specifically performing the experiments, or data/evidence collection | N/A |
| Resources | Provision of study materials, reagents, materials, patients, laboratory samples, animals, instrumentation, computing resources, or other analysis tools | TS, MK, JA |
| Data Curation | Management activities to annotate (produce metadata), scrub data and maintain research data (including software code, where it is necessary for interpreting the data itself) for initial use and later reuse | MK, JA |
| Writing - Original Draft | Preparation, creation and/or presentation of the published work, specifically writing the initial draft (including substantive translation) | MK, NSW |
| Writing - Review & Editing | Preparation, creation and/or presentation of the published work by those from the original research group, specifically critical review, commentary or revision – including pre-or postpublication stages | MK, TS, CEM, JA, NSW |
| Visualization | Preparation, creation and/or presentation of the published work, specifically visualization/ data presentation | MK, NSW |
| Supervision | Oversight and leadership responsibility for the research activity planning and execution, including mentorship external to the core team | NSW |
| Project administration | Management and coordination responsibility for the research activity planning and execution | MK |
| Funding acquisition | Acquisition of the financial support for the project leading to this publication | NSW, CEM |

## Notes

### Competing Interest Statement

The authors have declared no competing interest.

## References

1. Hamilton WD. The genetical evolution of social behaviour. I. Journal of theoretical biology. 1964;7(1):1–16.

2. West SA, Griffin AS, Gardner A. Evolutionary explanations for cooperation. Curr Biol. 2007;17(16):R661–72.

3. Strassmann JE, Gilbert OM, Queller DC. Kin discrimination and cooperation in microbes. Annu Rev Microbiol. 2011;65:349–67.

4. Ho HI, Hirose S, Kuspa A, Shaulsky G. Kin recognition protects cooperators against cheaters. Curr Biol. 2013;23(16):1590–5.

5. Strassmann JE, Queller DC. Evolution of cooperation and control of cheating in a social microbe. Proc Natl Acad Sci USA. 2011;108 Suppl 2(Suppl 2):10855–62.

6. Stefanic P, Belcijan K, Kraigher B, Kostanjsek R, Nesme J, Madsen JS, et al. Kin discrimination promotes horizontal gene transfer between unrelated strains in Bacillus subtilis. Nature Communications. 2021;12(1).

7. Stefanic P, Kraigher B, Lyons NA, Kolter R, Mandic-Mulec I. Kin discrimination between sympatric Bacillus subtilis isolates. Proc Natl Acad Sci USA. 2015;112(45):14042–7.

8. Lyons NA, Kraigher B, Stefanic P, Mandic-Mulec I, Kolter R. A Combinatorial Kin Discrimination System in Bacillus subtilis. Curr Biol. 2016;26(6):733–42.

9. Kearns DB. A field guide to bacterial swarming motility. Nat Rev Microbiol. 2010;8(9):634–44.

10. Sansinenea E, Ortiz A. Secondary metabolites of soil Bacillus spp. Biotechnology letters. 2011;33(8):1523–38.

11. Kiesewalter HT, Lozano-Andrade CN, Wibowo M, Strube ML, Maroti G, Snyder D, et al. Genomic and Chemical Diversity of Bacillus subtilis Secondary Metabolites against Plant Pathogenic Fungi. mSystems. 2021;6(1).

12. Eigentler L, Kalamara M, Ball G, MacPhee CE, Stanley-Wall NR, Davidson FA. Founder cell configuration drives competitive outcome within colony biofilms. ISME J. 2022.

13. Eigentler L, Davidson FA, Stanley-Wall NR. Mechanisms driving spatial distribution of residents in colony biofilms: an interdisciplinary perspective. Open Biol. 2022;12(12):220194.

14. Kobayashi K. Diverse LXG toxin and antitoxin systems specifically mediate intraspecies competition in Bacillus subtilis biofilms. PLoS Genet. 2021;17(7):e1009682.

15. Harwood CR, Cutting SM. Molecular biological methods for Bacillus. John Wiley & Sons Ltd. Chichester, England. 1990.

16. Verhamme DT, Kiley TB, Stanley-Wall NR. DegU co-ordinates multicellular behaviour exhibited by Bacillus subtilis. Mol Microbiol. 2007;65(2):554–68.

17. Konkol MA, Blair KM, Kearns DB. Plasmid-encoded Coml inhibits competence in the ancestral strain of Bacillus subtilis. Journal of Bacteriology. 2013.

18. Kalamara M, Abbott JC, MacPhee CE, Stanley-Wall NR. Biofilm hydrophobicity in environmental isolates of Bacillus subtilis. Microbiology (Reading). 2021;167(9).

19. Stanley NR, Britton RA, Grossman AD, Lazazzera BA. Identification of catabolite repression as a physiological regulator of biofilm formation by *Bacillus subtilis* by use of DNA microarrays. J Bacteriol. 2003;185(6):1951–7.

20. Gillespie RM, Stanley-Wall NR. Enzymes in action: an interactive activity designed to highlight positive attributes of extracellular enzymes synthesized by microbes. Journal of microbiology & biology education. 2014;15(2):310–2.

21. Allan C, Burel JM, Moore J, Blackburn C, Linkert M, Loynton S, et al. OMERO: flexible, model-driven data management for experimental biology. Nat Methods. 2012;9(3):245–53.

22. Bolger AM, Lohse M, Usadel B. Trimmomatic: a flexible trimmer for Illumina sequence data. Bioinformatics. 2014;30(15):2114–20.

23. Kolmogorov M, Yuan J, Lin Y, Pevzner PA. Assembly of long, error-prone reads using repeat graphs. Nat Biotechnol. 2019;37(5):540–6.

24. Wick RR, Judd LM, Gorrie CL, Holt KE. Unicycler: Resolving bacterial genome assemblies from short and long sequencing reads. PLoS Comput Biol. 2017;13(6):e1005595.

25. Schwengers O, Jelonek L, Dieckmann MA, Beyvers S, Blom J, Goesmann A. Bakta: rapid and standardized annotation of bacterial genomes via alignment-free sequence identification. Microb Genom. 2021;7(11).

26. Carver T, Harris SR, Berriman M, Parkhill J, McQuillan JA. Artemis: an integrated platform for visualization and analysis of high-throughput sequence-based experimental data. Bioinformatics. 2012;28(4):464–9.

27. Waterhouse AM, Procter JB, Martin DM, Clamp M, Barton GJ. Jalview Version 2--a multiple sequence alignment editor and analysis workbench. Bioinformatics. 2009;25(9):1189–91.

28. Kumar S, Stecher G, Tamura K. MEGA7: Molecular Evolutionary Genetics Analysis Version 7.0 for Bigger Datasets. Mol Biol Evol. 2016;33(7):1870–4.

29. Blin K, Shaw S, Kloosterman AM, Charlop-Powers Z, van Wezel GP, Medema MH, et al. antiSMASH 6.0: improving cluster detection and comparison capabilities. Nucleic Acids Res. 2021.

30. Lyons NA, Kolter R. Bacillus subtilis Protects Public Goods by Extending Kin Discrimination to Closely Related Species. MBio. 2017;8(4).

31. Kraigher B, Butolen M, Stefanic P, Mandic Mulec I. Kin discrimination drives territorial exclusion during Bacillus subtilis swarming and restrains exploitation of surfactin. ISME J. 2022;16(3):833–41.

32. Kalamara M, Spacapan M, Mandic-Mulec I, Stanley-Wall NR. Social behaviours by Bacillus subtilis: quorum sensing, kin discrimination and beyond. Mol Microbiol. 2018;110(6):863–78.

33. Page AJ, Cummins CA, Hunt M, Wong VK, Reuter S, Holden MT, et al. Roary: rapid large-scale prokaryote pan genome analysis. Bioinformatics. 2015;31(22):3691–3.

34. Patel PS, Huang S, Fisher S, Pirnik D, Aklonis C, Dean L, et al. Bacillaene, a novel inhibitor of procaryotic protein synthesis produced by Bacillus subtilis: production, taxonomy, isolation, physico-chemical characterization and biological activity. J Antibiot (Tokyo). 1995;48(9):997–1003.

35. Umezawa H, Aoyagi T, Nishikiori T, Okuyama A, Yamagishi Y, Hamada M, et al. Plipastatins: new inhibitors of phospholipase A2, produced by Bacillus cereus BMG302-fF67. I. Taxonomy, production, isolation and preliminary characterization. J Antibiot (Tokyo). 1986;39(6):737–44.

36. May JJ, Wendrich TM, Marahiel MA. The dhb operon of Bacillus subtilis encodes the biosynthetic template for the catecholic siderophore 2,3-dihydroxybenzoate-glycine-threonine trimeric ester bacillibactin. Journal of Biological Chemistry. 2001;276(10): 7209–17.

37. Arima K, Kakinuma A, Tamura G. Surfactin, a crystalline peptidelipid surfactant produced by *Bacillus subtilis:* isolation, characterization and its inhibition of fibrin clot formation. Biochemical and biophysical research communications. 1968;31(3):488–94.

38. Babasaki K, Takao T, Shimonishi Y, Kurahashi K. Subtilosin A, a new antibiotic peptide produced by Bacillus subtilis 168: isolation, structural analysis, and biogenesis. J Biochem. 1985;98(3):585–603.

39. Kenig M, Abraham EP. Antimicrobial activities and antagonists of bacilysin and anticapsin. Journal of general microbiology. 1976;94(1):37–45.

40. Phelan RW, Barret M, Cotter PD, O’Connor PM, Chen R, Morrissey JP, et al. Subtilomycin: a new lantibiotic from Bacillus subtilis strain MMA7 isolated from the marine sponge Haliclona simulans. Mar Drugs. 2013;11(6):1878–98.

41. Allenby NE, Watts CA, Homuth G, Pragai Z, Wipat A, Ward AC, et al. Phosphate starvation induces the sporulation killing factor of Bacillus subtilis. J Bacteriol. 2006; 188(14):5299–303.

42. Fawcett P, Eichenberger P, Losick R, Youngman P. The transcriptional profile of early to middle sporulation in *Bacillus subtilis*. Proceedings National Academy Sciences USA. 2000;97(14):8063–8.

43. Molle V, Fujita M, Jensen ST, Eichenberger P, Gonzalez-Pastor JE, Liu JS, et al. The Spo0A regulon of *Bacillus subtilis*. Mol Microbiol. 2003;50(5):1683–701.

44. Popp PF, Friebel L, Benjdia A, Guillot A, Berteau O, Mascher T. The Epipeptide Biosynthesis Locus epeXEPAB Is Widely Distributed in Firmicutes and Triggers Intrinsic Cell Envelope Stress. Microb Physiol. 2021.

45. Popp PF, Benjdia A, Strahl H, Berteau O, Mascher T. The Epipeptide YydF Intrinsically Triggers the Cell Envelope Stress Response of Bacillus subtilis and Causes Severe Membrane Perturbations. Frontiers in microbiology. 2020;11:151.

46. Paik SH, Chakicheria A, Hansen JN. Identification and characterization of the structural and transporter genes for, and the chemical and biological properties of, sublancin 168, a novel lantibiotic produced by Bacillus subtilis 168. Journal of Biological Chemistry. 1998;273(36):23134–42.

47. Dorenbos R, Stein T, Kabel J, Bruand C, Bolhuis A, Bron S, et al. Thiol-disulfide oxidoreductases are essential for the production of the lantibiotic sublancin 168. J Biol Chem. 2002;277(19):16682–8.

48. Butcher BG, Lin YP, Heimann JD. The yydFGHIJ operon of Bacillus subtilis encodes a peptide that induces the LiaRS two-component system. J Bacteriol. 2007;189(23):8616–25.

49. Benjdia A, Guillot A, Ruffie P, Leprince J, Berteau O. Post-translational modification of ribosomally synthesized peptides by a radical SAM epimerase in Bacillus subtilis. Nat Chem. 2017;9(7):698–707.

50. Hamon MA, Lazazzere BA. The sporulation transcription factor Spo0A is required for biofilm development in *Bacillus subtilis*. Mol Microbiol.2001;42(5):1199–209.

51. Kearns DB, Chu F, Brenda SS, Kolter R, Losick R. A master regulator for biofilm formation by *Bacillus subtilis*. Mol Microbiol. 2005;55(3):739–49.

