## Supplemental Information for "The role of the epipeptide EpeX in defining competitive fitness in intra-species mixed isolate colony biofilms of *Bacillus subtilis*"

*** For contact:**

Prof Nicola Stanley-Wall



**Keywords:** *Bacillus subtilis*, biofilm, kin discrimination, epipeptide, EpeX

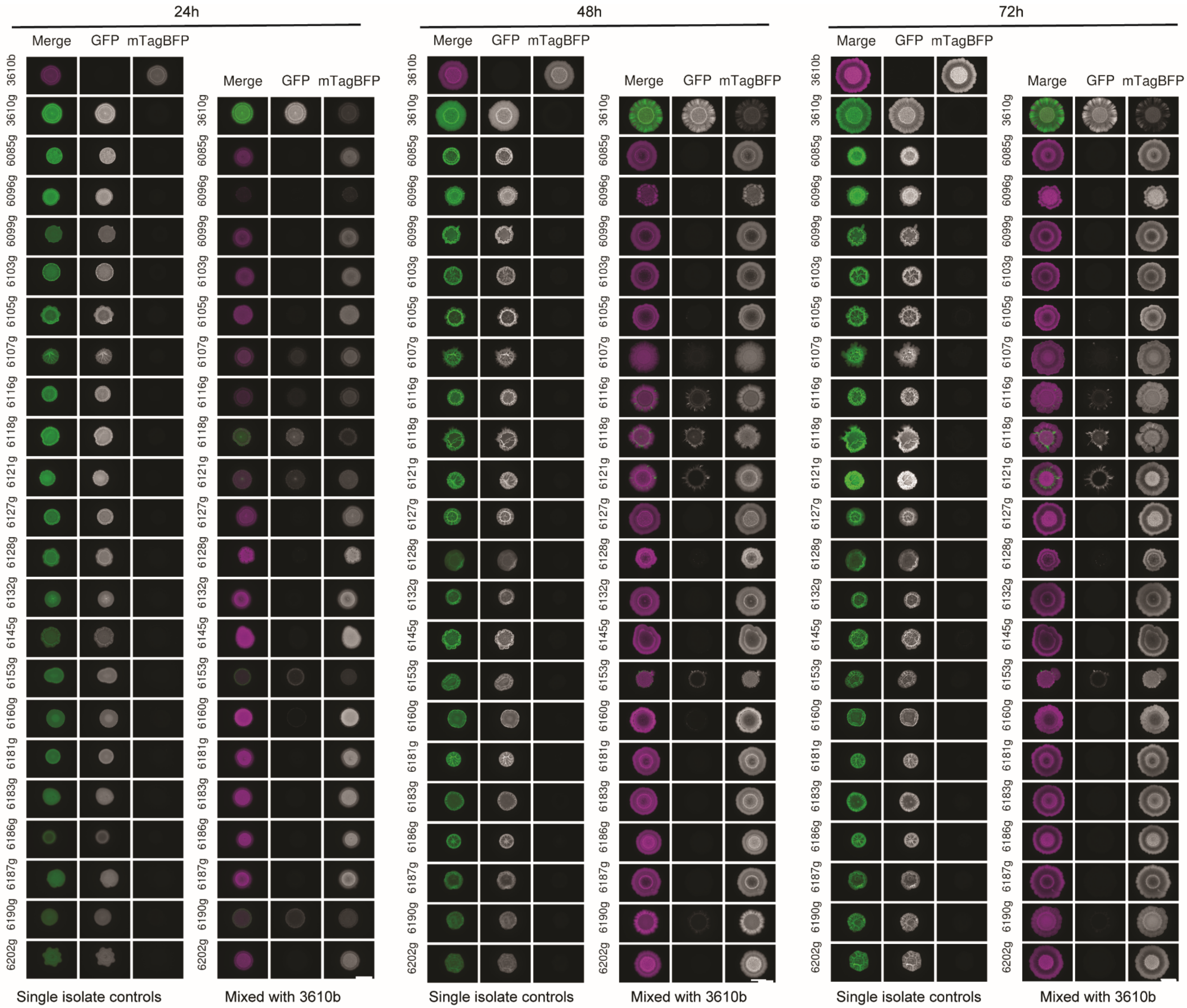
24h
48h
72h
Merge
GFP
mTagBFP
Marge
3610b
3610g
6085g
6096g
6099g
6103g
6105g
6107g
6116g
6118g
6121g
6127g
6128g
6132g
6145g
6153g
6160g
6181g
6183g
6186g
6187g
6190g
6202g
Single isolate controls
Mixed with 3610b

**Figure S1: Mixed isolate biofilm screen with NCIB 3610 as a reference.** Representative images from the mixed biofilm screen. Single isolate controls and mixed biofilms using NCIB 3610 mTagBFP as a reference are shown, taken after 24, 48 and 72h of growth as indicated. 3610 represents the model isolate NCIB 3610 and other numbers represent NRS numbers given to soil isolates in our collection (See Table 1). Isolate names followed by “g” represent strains constitutively expressing GFP, false coloured in green here, and isolate names followed by “b” represent strains expressing mTagBFP, false coloured in magenta. The labels at the top of each vertical panel represent the fluorescent channel images shown where “merge” shows both the GFP and mTagBFP channels. The scale bars represent 0.5 cm.

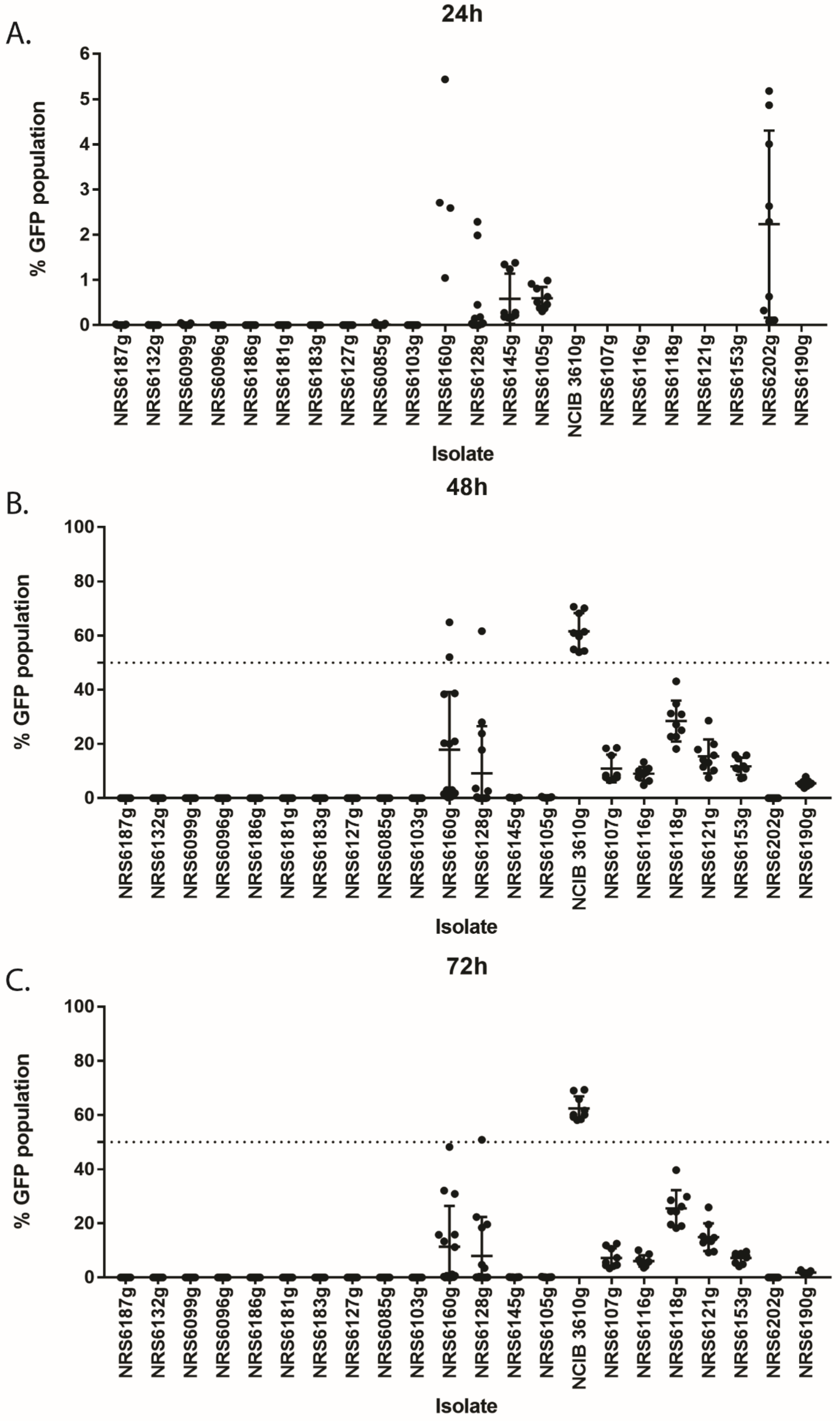

A.
24h
% GFP population
Isolate
B.
48h
% GFP population
Isolate
C.
72h
% GFP population
Isolate
NRS6187g
NRS6132g
NRS6099g
NRS6096g
NRS6186g
NRS6181g
NRS6183g
NRS6127g
NRS6085g
NRS6103g
NRS6160g
NRS6128g
NRS6145g
NRS6105g
NCIB 3610g
NRS6107g
NRS6116g
NRS6118g
NRS6121g
NRS6153g
NRS6202g
NRS6190g

**Figure S2: Outcome of biofilm co-culture screen with NCIB 3610 as a reference.** Competition results of NCIB 3610 mTagBFP against GFP-expressing soil isolates at 24 (A), 48 (B) and 72 (C) hours of co-incubation on biofilm-inducing media plates. The presented values are the % of the community of GFP expressing soil isolates, quantified using image analysis. The nine data points presented for each isolate represent three biological repeats and three technical repeats. The error bars represent the standard deviation of the mean. Note that data points in (A) are shown on a smaller scale for clarity.

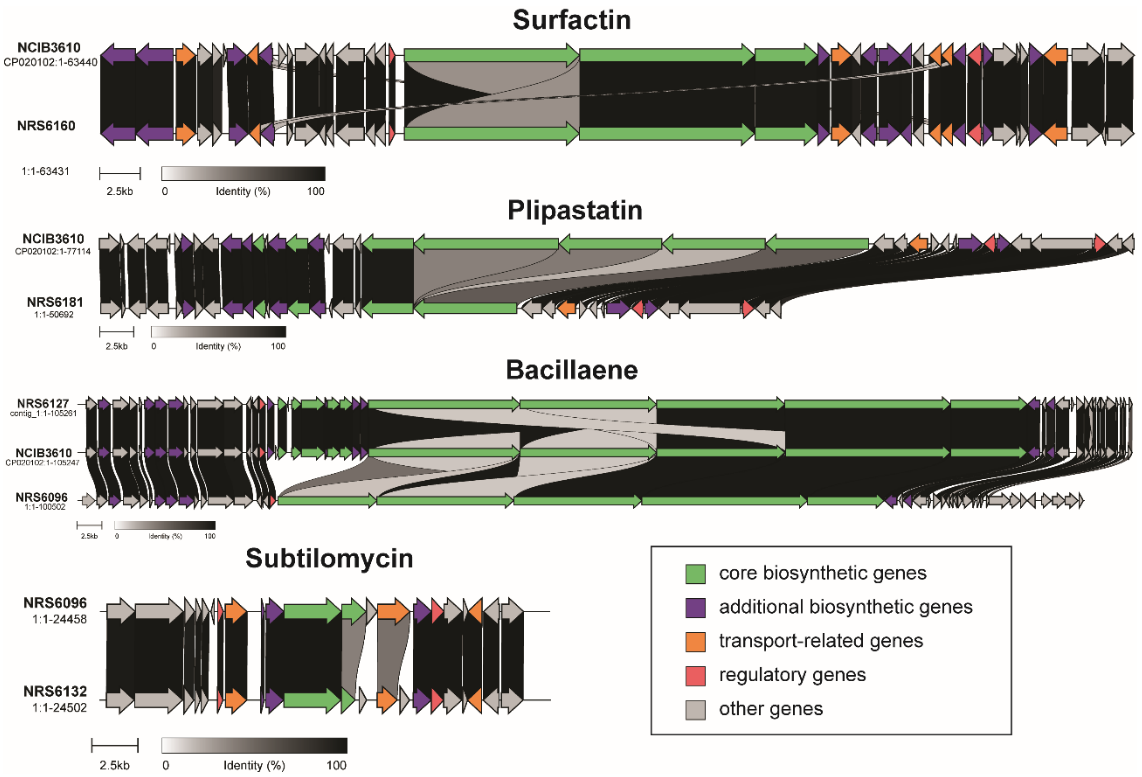


**Figure S3: Schematic representation of clusters found in *B. subtilis* genomes.** Shown are clusters that have some level of sequence variability from the consensus sequence of each of these clusters found within our isolate collection. In all cases, an isolate showing the default cluster is at the top with the variant cluster(s) found at the bottom. The diagrams were generated using clinker (1), using gbk files of the clusters produced by antiSMASH 6.0 (2) as input. The colour-coding annotation of the genes is described in the legend in the bottom right corner. The annotations presented here are based on the predicted function of each of these genes by antiSMASH.

## A.

1 M K I L K Y E D E K Y E V L V Q N N V F
1 ATGAAAATTTTAAAGTATGAAGATGAAAAATATGAAGTGCTTGTGCAAAACAACGTATTT

21 I K D K K S G E Y Y K N S L N S L S D K
61 ATAAAAGAC**AAAAAAA**GCGGAGAATACTATAAAAATAGTTTAAATTCTCTGTCAGATAAA

41 Q L L R F K M Y K E K V S P K F F Y L F
121 CAATTATTACGTTTTAAAATGTATAAAGAGAAAGTATCACCAAAGTTTTTTTATCTGTTC

61 L S F T A L M F I L N Y I H L I K L Q N
181 TTATCCTTTACGGCTCTCATGTTCATATTAAATTATATTCATTTAATAAAATTGCAAAAT

81 G L S S V F Y G W K M W I I I V I Y F I
241 GGATTGTCCTCAGTATTTTACGGCTGGAAAATGTGGATAATTATAGTTATTTATTTTATA

101 M N I V L H E L G H I Y S L K F F G K N
301 ATGAATATAGTGCTTCATGAACTGGGACATATTTATTCCTTAAAATTTTTCGGGAAAAAC

121 F D K V G F K L N F Y V F P A F Y V Q L
361 TTTGATAAAGTGGGATTCAAACTGAATTTTTATGTATTTCCTGCTTTTTATGTCCAATTG

141 N E T Y M L S R N E K I I V H L F G L F
421 AATGAAACCTACATGCTATCAAGAAATGAAAAGATAATCGTTCATTTATTTGGCTTGTTT

161 I N Y L L I N T L E L I N Q F T F S S E
481 ATCAATTATCTTTTAATAAATACATTAGAATTAATAAATCAATTTACATTTTCCAGTGAA

181 A L T M A F M L F S S T L L W N L I P I
541 GCATTAACTATGGCATTTATGCTTTTCTCGTCTACTTTATTGTGGAACTTAATCCCAATA

201 L N S D G Y K I L L A F L S L D E Y S R
601 TTGAATTCTGATGGTTACAAAATTCTTTTAGCTTTTCTGAGCTTAGATGAATATAGCCGT

221 F K T N H W L V L T I Q I I G I G L A V
661 TTTAAGACAAATCATTGGTTAGTTCTTACGATTCAAATTATAGGGATTGGCCTTGCAGTG

241 N S V V H W I L Y I V N *
721 AATTCAGTTGTACATTGGATATTATATATTGTAAACTAG

## B.

1 M K I L K Y E D E K Y E V L V Q N N V F
1 ATGAAAATTTTAAAGTATGAAGATGAAAAATATGAAGTGCTTGTGCAAAACAACGTATTT

21 I K D K K K A E N T I K I V *
61 ATAAAAGAC**AAAAAAAAA**GCGGAGAATACTATAAAAATAGTTTAAATTCTC...

**Figure S4: Nucleotide and corresponding amino acid sequences of *epeP* in isolates of *B. subtilis*.** (A) shows the nucleotide (bottom line) and corresponding amino acid (top line) sequences in isolates NCIB 3610, NRS6116, NRS6118, NRS6153, NRS6190 and NRS6202 and (B) shows the same sequences of isolates NRS6107 and NRS6121. The point at which the protein diverges is indicated in red. The asterisk represents a stop codon.

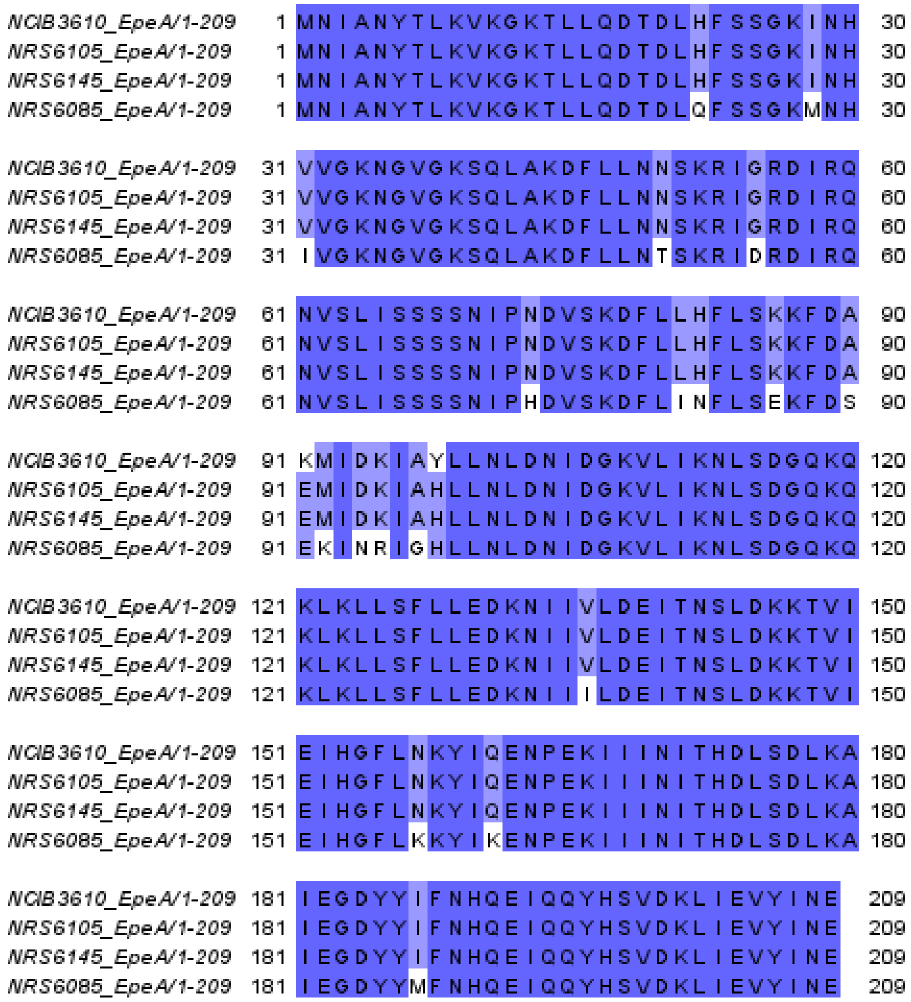


**Figure S5:** Alignment of EpeA sequences from *B. subtilis* isolates. Sequence alignments were performed in Jalview (3) using Mafft with default settings. The colouring scheme used represents percent identity, where amino acids coloured in dark blue correspond to an identity of >80% to the consensus sequence. The lighter shade of blue represents sequence identities of >60%. Amino acids coloured in white show a sequence identity of less than 40% to that of the consensus sequence.

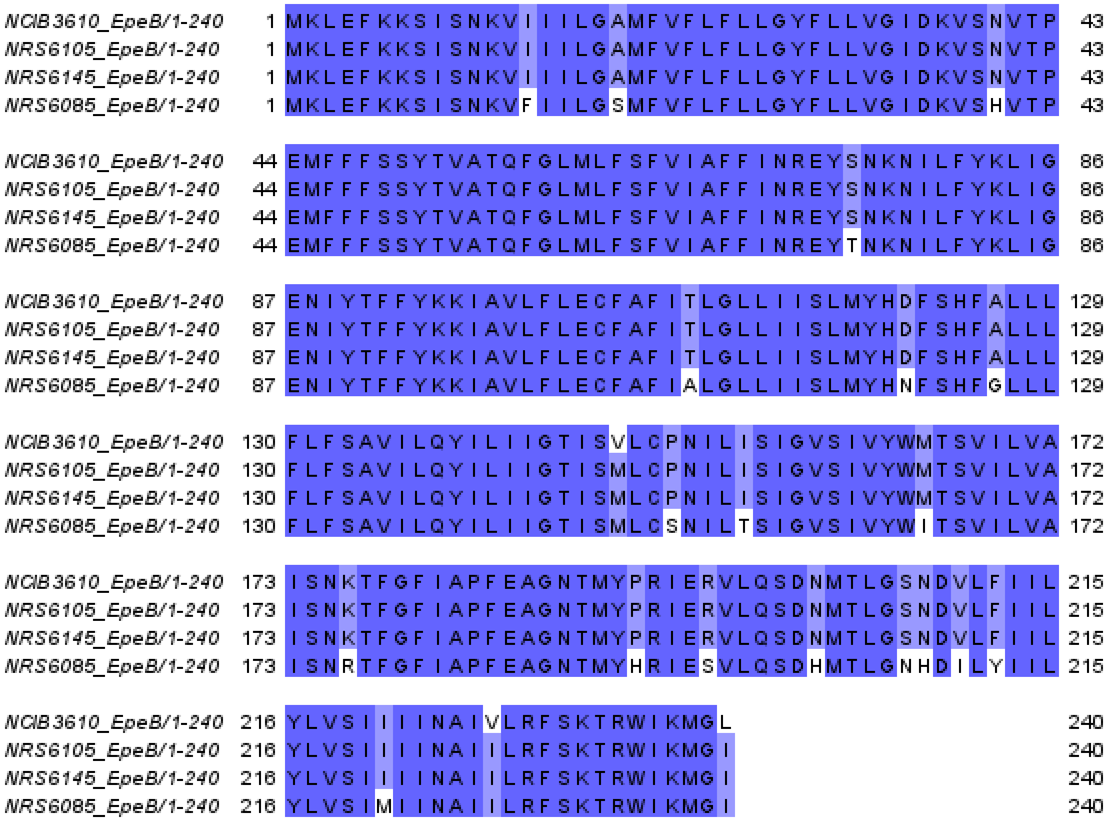


**Figure S6: Alignment of EpeB sequences from *B. subtilis* isolates.** Sequence alignments were performed in Jalview (3) using Mafft with default settings. The colouring scheme used represents percent identity, where amino acids coloured in dark blue correspond to an identity of >80% to the consensus sequence. The lighter shade of blue represents sequence identities of >60%. Amino acids coloured in white show a sequence identity of less than 40% to that of the consensus sequence.

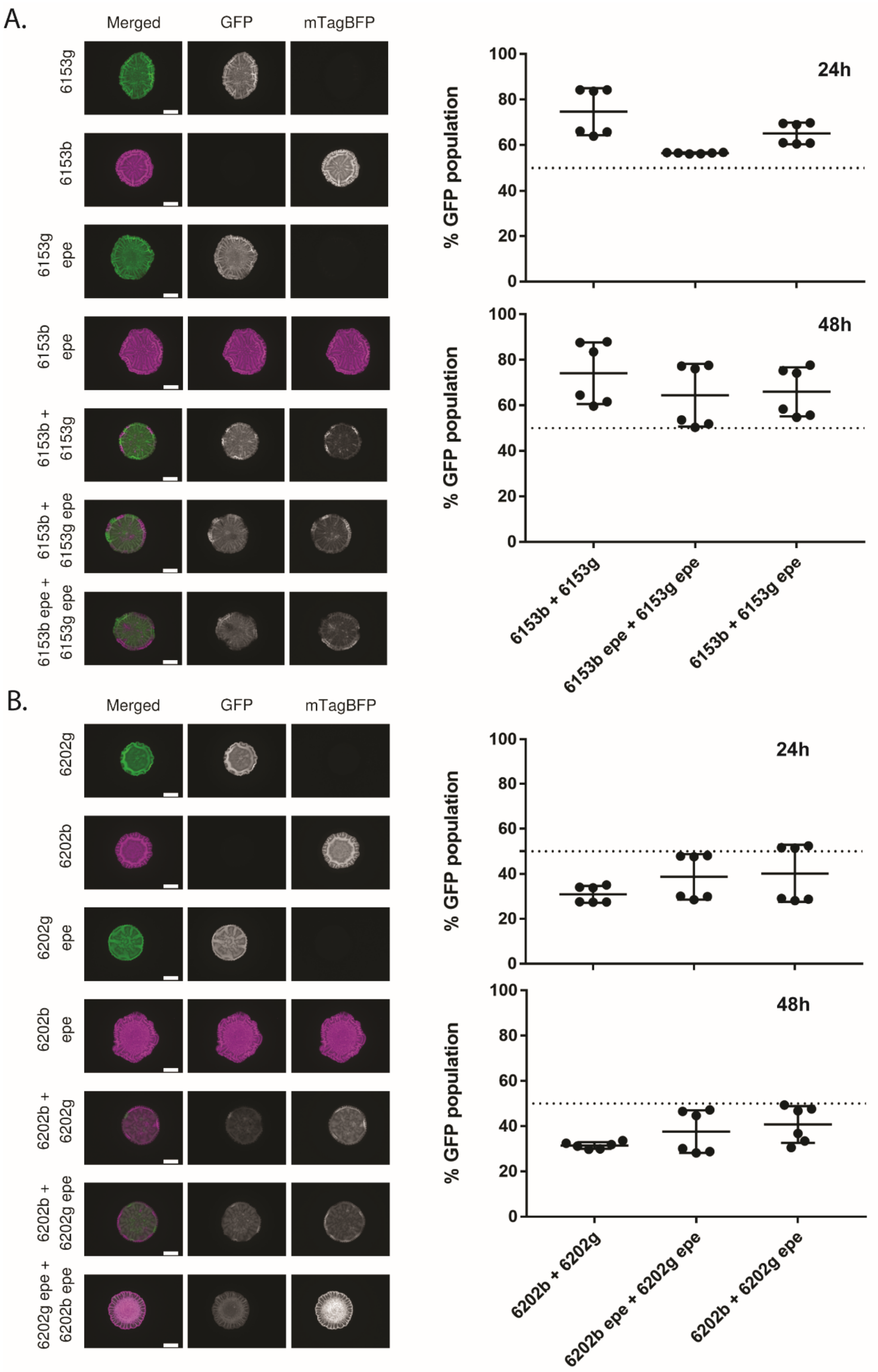

A.
Merged
GFP
mTagBFP
6153g
6153b
6153g epe
6153b epe
6153b + 6153g
6153b + 6153g epe
6153b epe + 6153g epe
% GFP population
24h
48h
0
20
40
60
80
100
6153b + 6153g
6153b epe + 6153g epe
6153b + 6153g epe
B.
Merged
GFP
mTagBFP
6202g
6202b
6202g epe
6202b epe
6202b + 6202g
6202b + 6202g epe
6202g epe + 6202b epe
% GFP population
24h
48h
0
20
40
60
80
100
6202b + 6202g
6202b epe + 6202g epe
6202b + 6202g epe

**Figure S7: Competition outcomes of *epeXEPAB* deletion strains of NRS6153 and NRS6202.** On the left, representative images of biofilms growth for 48 hours at 30°C on MSgg agar are shown. “6153” is the soil isolate NRS6153 (A) and “6202” is the isolate NRS6202 (B). Strain names followed by “b” represent strains constitutively expressing mTagBFP and names followed by “g” represent strains constitutively expressing GFP. “epe” represents deletion of the *epeXEPAB* operon. The scale bars represent 0.5 cm. On the right, competition results of either NRS6153 (A) or NRS6202 (B) wild type expressing mTagBFP (NRS6938 and NRS7201 respectively) against GFP-expressing wild type (NRS6222 and NRS6897) or *epeXEPAB* mutant (NRS7390 and NRS7392 respectively) of the soil isolates after 24 and 48 of co-incubation on biofilm inducing media plates as indicated. The presented values are the % of the community of GFP expressing strain, quantified using image analysis. Each individual data point presented for each isolate represent one of three technical replicates for the two biological repeats performed. The error bars represent the standard deviation of the mean.

**Table S1: Synthetic sequence used to construct the *epeXEPAB* mutants**

| plasmid | Construct sequence |
|---|---|
| pNW2315[a] | GGATCCGCGAGTAGATCTAGGTACTTGCTTTCCAAAACCGTCCATCCCCTTTTTAGAATAAACCCCTCTT<br>TATGAGCATGTGCATGATAAGGTGAATTTAAAAGTATGACATTATTTTTAAACATTGGACCTTTATATG<br>ATAGTACCGCATAACTGCTTTTAGAGACAATTAAAACGAGAGAAAGAGGTGCCAAATGGGGTAAGCA<br>AGAGCTTAAAACAGCTTTTCAGTACAGGTTGGTTTAGGGTAAGATTGTTTGCCTCATTTCTTCCACTTA<br>AAATCTGTATATCCCTTAACTAGTGTAGTCAATATTGATGTTTTACTTTTCATTACGAAGCTTTAAATCA<br>AACTATGGCCTTTCATTCGGTATAGCTTAGAGGGGCTAGAGGACTATAGGCTTTATATACCAAGAATA<br>GTGACATTTTTCAGATTTTTTAGGTACAATATATTGACATGTATTGAATGATATAGAATAATTGGTTTAT<br>ATTAGAAAAGGAGGAGGGATAATATGGTTTGATTTTTAATGGATAATGTGATATAATCTTTAAATACT<br>GTAGAAAAGAGGAAGGAAATAATAAATGGCTAAAATGAGAATATCACCGGAATTGAAAAAACTGATC<br>GAAAAATACCGCTGCGTAAAAGATACGGAAGGAATGTCTCCTGCTAAGGTATATAAGCTGGTGGGAG<br>AAAATGAAAACCTATATTTAAAAATGACGGACAGCCGGTATAAAGGGACCACCTATGATGTGGAACG<br>GGAAAAGGACATGATGCTATGGCTGGAAGGAAAGCTGCCTGTTCCAAAGGTCCTGCACTTTGAACGG<br>CATGATGGCTGGAGCAATCTGCTCATGAGTGAGGCCGATGGCGTCCTTTGCTCGGAAGAGTATGAAG<br>ATGAACAAAGCCCTGAAAAGATTATCGAGCTGTATGCGGAGTGCATCAGGCTCTTTCACTCCATCGAC<br>ATATCGGATTGTCCCTATACGAATAGCTTAGACAGCCGCTTAGCCGAATTGGATTACTTACTGAATAAC<br>GATCTGGCCGATGTGGATTGCGAAAACTGGGAAGAAGACACTCCATTTAAAGATCCGCGCGAGCTGT<br>ATGATTTTTTAAAGACGGAAAAGCCCGAAGAGGAACTTGTCTTTTCCCACGGCGACCTGGGAGACAG<br>CAACATCTTTGTGAAAGATGGCAAAGTAAGTGGCTTTATTGATCTTGGGAGAAGCGGCAGGGCGGAC<br>AAGTGGTATGACATTGCCTTCTGCGTCCGGTCGATCAGGGAGGATATCGGGGAAGAACAGTATGTCG<br>AGCTATTTTTTGACTTACTGGGGATCAAGCCTGATTGGGAGAAAATAAAATATTATATTTTACTGGATG<br>AATTGTTTTAGTACCTAGATAATAAAGGGAGCTACTTCACAATGAAGTAGCTTTTTATGTGCACCTATA<br>TATCATTAACATCAAGATTAATGCATGTGACTGGCTGAGTTTGTTAAGGGAAGGATGTTTTTCTGTTTG<br>CATCGCCCAAACTATTATTGTTTTTCTTTCACCATCTTTACAAAGAGGAAAAAGATAATGAATACGATA<br>AAGTAGTTCATTGGATTCTTAACGATGCTCATCTCTAGTGAAATCTGTAAGAGTATCCAGAGCGCAAA<br>GATGGCTAGTATGGCAGGAATGATAATTTTGTACATGTCGACCACCTACCGTAGTGAAGAGCCCTATA<br>ATAAAAAACTATTAGCCTGTACCACAAACTGCGACTGCTCATAATTATACGAAATCTTCGAGTAATCAA<br>ACGGCTGCCCATTTGTCAGGTGAAAAATACTTTCTATATACAGCTTCGGCAGTCCAGCCTCTAATCCCA<br>AATATTCCGCTTCTTCCTCATTAAGCTGGCCCACGTGCCTGCAG |

.

[a] nucleotides 7 to 509 correspond to base pairs 4127683 to 4128185 of the NCIB 3610 genome; nucleotides 1,380 to 1,879 are base pairs 4123761 to 4124260 of the NCIB 3610 genome; and nucleotides 510 to 1,379 carry the kanamycin resistance cassette. The construct was inserted into a pUC57 cassette using BamHI/PstI restriction sites.

**Table S2: Overview of assembled genomes of *B. subtilis***

| Isolate | Accession | Chromosomal Contigs | Plasmids | Length (bp) | N50 (bp) | GC (%) | Annotated CDS | Annotated rRNA | Annotated tRNA |
|---|---|---|---|---|---|---|---|---|---|
| NRS6085 | ERZ15916258 | 1 | 1 | 4,342,529 | 4,335,289 | 43.4 | 4,405 | 30 | 86 |
| NRS6096 | ERZ15916259 | 2 | 3 | 4,312,183 | 4,128,244 | 43.4 | 4,330 | 30 | 88 |
| NRS6099 | ERZ15916260 | 1 | 0 | 4,096,107 | 4,096,107 | 43.8 | 4,064 | 30 | 86 |
| NRS6103 | ERZ15916261 | 1 | 0 | 4,264,553 | 4,264,553 | 43.5 | 4,271 | 30 | 86 |
| NRS6105 | ERZ15916262 | 1 | 0 | 4,090,679 | 4,090,679 | 43.8 | 4,021 | 30 | 86 |
| NRS6107 | ERZ15916263 | 1 | 1 | 4,166,364 | 4,101,676 | 43.7 | 4,126 | 30 | 86 |
| NRS6116 | ERZ15916264 | 1 | 1 | 4,371,489 | 4,307,899 | 43.3 | 4,448 | 27 | 86 |
| NRS6118 | ERZ15916265 | 1 | 0 | 4,261,356 | 4,261,356 | 43.5 | 4,282 | 30 | 86 |
| NRS6121 | ERZ15916267 | 1 | 0 | 4,123,859 | 4,123,859 | 43.8 | 4,084 | 30 | 86 |
| NRS6127 | ERZ15916268 | 1 | 0 | 4,226,657 | 4,226,657 | 43.5 | 4,244 | 30 | 86 |
| NRS6128 | ERZ15916269 | 1 | 0 | 4,001,615 | 4,001,615 | 43.9 | 3,930 | 30 | 86 |
| NRS6131 | ERZ15916270 | 1 | 1 | 4,257,224 | 4,249,983 | 43.2 | 4,334 | 30 | 87 |
| NRS6132 | ERZ15916271 | 1 | 1 | 4,181,988 | 4,097,627 | 43.6 | 4,162 | 30 | 88 |
| NRS6145 | ERZ15916273 | 1 | 1 | 4,169,376 | 4,085,688 | 43.6 | 4,117 | 27 | 88 |
| NRS6153 | ERZ15916274 | 1 | 0 | 4,215,744 | 4,215,744 | 43.7 | 4,211 | 30 | 86 |
| NRS6160 | ERZ15916275 | 1 | 0 | 4,224,542 | 4,224,542 | 43.6 | 4,218 | 30 | 86 |
| NRS6167 | ERZ15916276 | 1 | 0 | 4,192,866 | 4,192,866 | 44 | 4,116 | 30 | 86 |
| NRS6181 | ERZ15916277 | 1 | 1 | 4,275,379 | 4,189,561 | 43.6 | 4,338 | 30 | 88 |
| NRS6183 | ERZ15916278 | 1 | 0 | 4,256,610 | 4,256,610 | 43.5 | 4,283 | 30 | 86 |
| NRS6186 | ERZ15916279 | 1 | 1 | 4,324,000 | 4,239,628 | 43.3 | 4,363 | 30 | 88 |
| NRS6187 | ERZ15916280 | 1 | 0 | 4,096,109 | 4,096,109 | 43.8 | 4,063 | 30 | 86 |
| NRS6190 | ERZ15916281 | 2 | 1 | 4,169,151 | 4,056,987 | 43.7 | 4,147 | 30 | 86 |
| NRS6202 | ERZ15916282 | 3 | 0 | 4,248,983 | 4,243,504 | 43.5 | 4,283 | 24 | 86 |

| Type of molecule | Most similar known cluster | % ID | Biosynthetic gene |
|---|---|---|---|
| Ranthipeptide, sactipeptide | Sporulation killing factor | 100% | *skfC* |
| Glycocin | Sublancin 168 | 100% | *sunA* |
| RRE-containing | | | *aslA* (*yxbA*) |
| CDPS | | | *yvmC* |
| Lanthipeptide | subtilomycin | 100% / 83% | *subB, subC* |
| Epipeptide | thailanstatin A | 10% | *yydG (epeE), yydH (epeX)* |

**Table S3: Accessory clusters predicted by antiSMASH v.6.0.** RRE stands for RiPP (ribosomally synthesized and posttranslationally modified peptide) recognition element. CDPS stands for tRNA-dependent cyclodipeptide synthases. % ID shows the percent identity of the cluster to the most similar known cluster in the AntiSMASH database and the biosynthetic gene is the core biosynthetic identified gene(s) in the *B. subtilis* genome. Gene names in brackets indicate synonyms of gene names. For the lanthipeptide subtilomycin, the % ID is presented is 100 for all isolates encoding the cluster apart from one (NRS6132), which has a % ID of 83 compared to the rest.

| Region | Isolate | % ID | alignment length (bp) | mismatches | gap opens | s. start | s. end | E value |
|---|---|---|---|---|---|---|---|---|
| *epeX* | NCIB 3610 | 100 | 150 | 0 | 0 | 4127685 | 4127536 | 2.10E-74 |
| *epeX* | NRS6107 | 100 | 150 | 0 | 0 | 4013218 | 4013069 | 2.10E-74 |
| *epeX* | NRS6116 | 100 | 150 | 0 | 0 | 4165587 | 4165438 | 2.10E-74 |
| *epeX* | NRS6118 | 100 | 150 | 0 | 0 | 1380910 | 1380761 | 2.10E-74 |
| *epeX* | NRS6121 | 100 | 150 | 0 | 0 | 270958 | 270809 | 2.10E-74 |
| *epeX* | NRS6153 | 100 | 150 | 0 | 0 | 4127374 | 4127225 | 2.10E-74 |
| *epeX* | NRS6190 | 100 | 150 | 0 | 0 | 4016655 | 4016506 | 2.10E-74 |
| *epeX* | NRS6202 | 100 | 150 | 0 | 0 | 4170601 | 4170452 | 2.10E-74 |
| *epeE* | NCIB 3610 | 100 | 960 | 0 | 0 | 4127478 | 4126519 | 0 |
| *epeE* | NRS6107 | 100 | 960 | 0 | 0 | 4013011 | 4012052 | 0 |
| *epeE* | NRS6116 | 100 | 960 | 0 | 0 | 4165380 | 4164421 | 0 |
| *epeE* | NRS6118 | 100 | 960 | 0 | 0 | 1380703 | 1379744 | 0 |
| *epeE* | NRS6121 | 100 | 960 | 0 | 0 | 270751 | 269792 | 0 |
| *epeE* | NRS6153 | 100 | 960 | 0 | 0 | 4127167 | 4126208 | 0 |
| *epeE* | NRS6190 | 100 | 960 | 0 | 0 | 4016448 | 4015489 | 0 |
| *epeE* | NRS6202 | 100 | 960 | 0 | 0 | 4170394 | 4169435 | 0 |
| *epeP* | NCIB 3610 | 100 | 759 | 0 | 0 | 4126538 | 4125780 | 0 |
| *epeP* | NRS6107 | 99.474 | 761 | 0 | 2 | 4012071 | 4011313 | 0 |
| *epeP* | NRS6116 | 100 | 759 | 0 | 0 | 4164440 | 4163682 | 0 |
| *epeP* | NRS6118 | 100 | 759 | 0 | 0 | 1379763 | 1379005 | 0 |
| *epeP* | NRS6121 | 99.215 | 764 | 0 | 3 | 269811 | 269049 | 0 |
| *epeP* | NRS6153 | 100 | 759 | 0 | 0 | 4126227 | 4125469 | 0 |
| *epeP* | NRS6190 | 100 | 759 | 0 | 0 | 4015508 | 4014750 | 0 |
| *epeP* | NRS6202 | 100 | 759 | 0 | 0 | 4169454 | 4168696 | 0 |
| *epeA* | NCIB3610 | 100 | 630 | 0 | 0 | 4125630 | 4125001 | 0 |

| Region | Isolate | % ID | alignment length (bp) | mismatches | gap opens | s. start | s. end | E value |
|---|---|---|---|---|---|---|---|---|
| *epeA* | NRS6107 | 100 | 630 | 0 | 0 | 4011163 | 4010534 | 0 |
| *epeA* | NRS6116 | 100 | 630 | 0 | 0 | 4163532 | 4162903 | 0 |
| *epeA* | NRS6118 | 100 | 630 | 0 | 0 | 1378855 | 1378226 | 0 |
| *epeA* | NRS6121 | 100 | 630 | 0 | 0 | 268899 | 268270 | 0 |
| *epeA* | NRS6153 | 100 | 630 | 0 | 0 | 4125319 | 4124690 | 0 |
| *epeA* | NRS6190 | 100 | 630 | 0 | 0 | 4014600 | 4013971 | 0 |
| *epeA* | NRS6202 | 100 | 630 | 0 | 0 | 4168546 | 4167917 | 0 |
| *epeA* | NRS6145 | 99.206 | 630 | 5 | 0 | 3990995 | 3990366 | 0 |
| *epeA* | NRS6105 | 99.206 | 630 | 5 | 0 | 3995988 | 3995359 | 0 |
| *epeA* | NRS6085 | 88.254 | 630 | 74 | 0 | 2301933 | 2302562 | 0 |
| *epeB* | NCIB 3610 | 100 | 723 | 0 | 0 | 4124980 | 4124258 | 0 |
| *epeB* | NRS6107 | 100 | 723 | 0 | 0 | 4010513 | 4009791 | 0 |
| *epeB* | NRS6116 | 100 | 723 | 0 | 0 | 4162882 | 4162160 | 0 |
| *epeB* | NRS6118 | 100 | 723 | 0 | 0 | 1378205 | 1377483 | 0 |
| *epeB* | NRS6121 | 100 | 723 | 0 | 0 | 268249 | 267527 | 0 |
| *epeB* | NRS6153 | 100 | 723 | 0 | 0 | 4124669 | 4123947 | 0 |
| *epeB* | NRS6190 | 100 | 723 | 0 | 0 | 4013950 | 4013228 | 0 |
| *epeB* | NRS6202 | 100 | 723 | 0 | 0 | 4167896 | 4167174 | 0 |
| *epeB* | NRS6145 | 99.032 | 723 | 1 | 0 | 3990345 | 3989623 | 0 |
| *epeB* | NRS6105 | 99.032 | 723 | 1 | 0 | 3995338 | 3994616 | 0 |
| *epeB* | NRS6085 | 87.866 | 717 | 87 | 0 | 2302583 | 2303299 | 0 |

**Table S4: Results of nucleotide BLAST analysis of the *epeX*, *epeE*, *epeP*, *epeA* and *epeB* coding sequences from *B. subtilis* 168 against the isolates used in this work.** "s." in "s. start" and "s. end" strands for "subject" and these values show the coordinate on the chromosome of each subject isolate where the indicated gene sequence starts and ends. % ID is the percent identity of the query sequence to the subject. The alignment length is in base pairs.

**Supplemental References**

1. Gilchrist CLM, Chooi YH. 2021. Clinker & clustermap.js: Automatic generation of gene cluster comparison figures. Bioinformatics doi:10.1093/bioinformatics/btab007.
2. Blin K, Shaw S, Kloosterman AM, Charlop-Powers Z, van Wezel GP, Medema MH, Weber T. 2021. antiSMASH 6.0: improving cluster detection and comparison capabilities. Nucleic Acids Res doi:10.1093/nar/gkab335.
3. Waterhouse AM, Procter JB, Martin DM, Clamp M, Barton GJ. 2009. Jalview Version 2--a multiple sequence alignment editor and analysis workbench. Bioinformatics 25:1189-91.
